# Self-confined expression in the *Arabidopsis* root stem cell niche

**DOI:** 10.1101/2021.12.02.468941

**Authors:** Josep Mercadal, Isabel Betegón-Putze, Nadja Bosch, Ana I. Caño-Delgado, Marta Ibañes

## Abstract

Stem cell niches are local microenvironments that preserve their unique identity while communicating with adjacent tissues. In the primary root of *Arabidopsis thaliana*, the stem cell niche comprises the expression of two transcription factors, BRAVO and WOX5, among others. Intriguingly, these proteins confine their own gene expression to the niche, as evidenced in each mutant background. Here we propose through mathematical modeling that BRAVO confines its own expression domain to the stem cell niche by attenuating its WOX5-dependent diffusible activator. This negative feedback drives WOX5 action to be spatially restricted as well. The results show that WOX5 diffusion and sequestration by binding to BRAVO is sufficient to drive realistic confined *BRAVO* expression at the stem cell niche. We propose that attenuation of a diffusible activator can be a general mechanism to confine genetic activity to a small region while at the same time maintain signaling within it and with the surrounding cells.

## 1 Introduction

Both in animals and plants, stem cells are maintained in tightly regulated microenvironments called stem cell niches (SCNs). Within these SCN, stem cells remain in an undifferentiated state and provide a continuous flux of precursors of more specialized cells that sustain growth and replace old or damaged tissues [1]. SCNs usually consist on a few number of stem cells maintained by short-range signals produced by localized sources or *organizing centers*, groups of cells which maintain neighboring cells in a stem cell state [2, 3]. As stem cell daughters are placed outside the reach of these signals, they begin to differentiate and give rise to more specialized cell types [4]. In animals, common signals preserving stem cells are diffusible ligands like the *Dpp* morphogen for *Drosophila* male or female germ cells [5, 6] and *Hedgehog* in mouse and *Drosophila* epithelial cells [7, 8], to name a few. Overproduction of these signals can drive an increase in the number of stem cells within the tissue, resulting in enlarged niches and often leading to malfunctioning of the surrounding tissue, or even whole organs [3]. Knowing the origin and function of these signals is therefore essential to understand the role of stem cells in the processes underlying organism development and sustenance.

In the model plant *Arabidopsis thaliana*, highly mobile hormones, such as auxin [9, 10, 11], as well as short-range moving transcription factors like *WUSCHEL* and *WOX5* (WUSCHEL-RELATED HOME-OBOX 5) [12, 13] are involved in specifying stem cell niches. In the root apical meristem of *Arabidopsis*, the SCN lies at the tip of the root, a location known to be established by positional information conferred by auxin signaling [14, 15]. In particular, the root SCN is specified by the overlapping of the SCARECROW (SCR) and SHORTROOT (SHR) transcription factors, together with the activation of PLETHORA (PLT) genes by the hormone auxin, whose levels peak at the position where the SCN is established [16]. This positional signaling allows for the necessary plasticity to establish a new niche when it has been destroyed or damaged, by virtue of a continual supply and renewal of stem cells at the very same location [17].

The root SCN is dynamically specified by the constant balance between external signaling and local communication, restricting the location of stem cells to a small, well-defined region. The SCN is formed by a small group of rarely dividing pluripotent cells called the quiescent center (QC) and by immediately surrounding stem cells, i.e. the vascular initials (VI), columella stem cells (CSC) and cortex-endodermis initials (CEI) (Figure 1A) [18]. Direct cell-cell contact from the QC to its surrounding stem cells is important for stem cell identity [19, 20] and can involve the transport of short-range signals from the QC. The homeodomain transcription factor WOX5 is specifically expressed at the QC [13] and is able to move towards adjacent cells [21, 22, 23]. WOX5 itself has been proposed to act as the long-sought short-range signal to repress columella stem cell differentiation [21], albeit recent results challenge this view [22]. While short-range signaling is thought to ensure that stem cell numbers are restrained and the SCN does not become displaced from the growing root tip, it is yet unclear how this is achieved [24, 25, 26].

**Figure 1:**
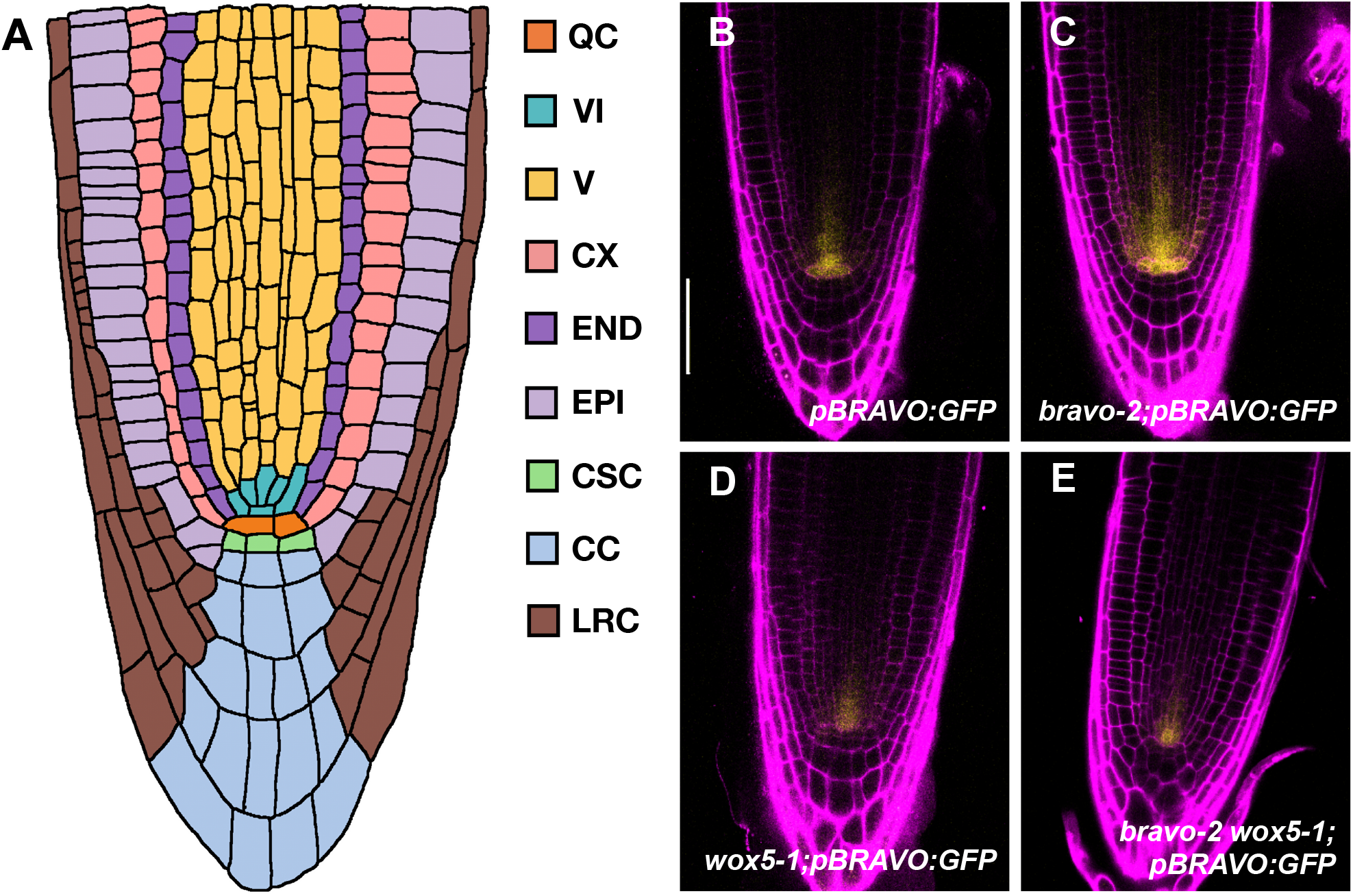
*pBRAVO-GFP* activity in WT and in loss of function mutants. **A)** Cartoon of the root apical meristem depicting its organization in cell-types: quiescent center (QC), vascular initials (V), vascular cells (V), cortex (CX), endodermis (END), epidermis (EPI), columella stem cells (CSC), columella cells (CC), lateral root cap (LRC). **B-E)** Confocal images of PI-stained 6-day-old roots. GFP-tagged expression is shown in yellow. Activity of the BRAVO promoter in WT (B), *bravo-2* (C), *wox5-1* (D) and *bravo-2 wox5-1* (E) loss-of-function backgrounds. Scale bar: 50*μ*m. The promoter activity of BRAVO expands its domain in the *bravo-2* mutant. This suggests that BRAVO confines its own expression to the QC and VI in the WT. The expansion is not observed in *wox5-1* mutants nor in *bravo-2 wox5-1* mutants, suggesting that such confinement requires WOX5. B-E from experiments in [28].

The R2R3-MYB transcription factor BRAVO (BRASSINOSTEROIDS AT VASCULAR AND OR-GANIZING CENTER) has recently been linked to the maintenance of SCN homeostasis [27, 28]. BRAVO is expressed at the QC and vascular initials [27], and colocalizes with WOX5 in the QC [28]. Both BRAVO and WOX5 have been shown to individually promote quiescence, as mutant roots with disrupted BRAVO or WOX5 exhibit increased QC divisions, supporting their role as essential factors for QC homeostasis [27, 29]. We have recently shown that BRAVO and WOX5 are codependent, as evidenced by the mutual regulation of each other promoter expression and the physical interaction of their corresponding proteins, presumably into a protein (e.g. heterodimer) complex [28]. These data also showed that the expression of the BRAVO promoter, restricted to the QC and VI in WT plants, expands towards the vasculature, cortex and endodermis in the *bravo* loss-of-function mutant [28] (Figure 1B,C), suggesting that BRAVO actively confines its own expression domain. Moreover, this expansion is not observed in the loss-of-function *wox5* mutant nor in the double loss-of-function mutant *bravo wox5* [28] (Figure 1D,E), pointing to a mechanism for self-confinement that is strongly WOX5-dependent. How this active confinement is achieved remains to be elucidated.

In this paper we show, through mathematical and computational analysis, that the spatial confinement of *BRAVO* expression can result from a negative feedback through a mobile activator, which might be WOX5 or a target thereof. This mechanism also reduces the spatial domain of WOX5 activity, overall providing a natural way for stem-specific factors to locally regulate SCN maintenance. We test different scenarios for the interactions between BRAVO and its activator, and study the implications and plausibility of each of them to explain the changes in expression observed experimentally. Our results support that the small diffusion of WOX5 [21, 22, 23], together with its physical interaction with BRAVO [28], can explain the confined nature of *BRAVO* expression. Additional interactions, which involve BRAVO and WOX5 regulating common targets in an antagonistic manner, and are supported by transcriptomic data on the QC [28], can act redundantly, but are required when WOX5 targets diffuse. Altogether, our results shed light on the regulatory principles balancing the confinement of transcription factors to a microenvironment and their communication with the surrounding cells.

## 2 Results

### 2.1 BRAVO can confine its own expression domain by immobilizing WOX5

The changes in the expression of *BRAVO* in the WT and in loss of function mutants of BRAVO and/or WOX5 [28] (Figure 1B-E) suggest that, in the WT, BRAVO confines its own expression to the SCN and that this occurs through a mechanism that requires WOX5. To decipher how this self-confinement can be attained, we first took into account that BRAVO transcription is ultimately activated by WOX5 (either directly or through WOX5-targets) [28] and that WOX5 proteins are able to move from the QC to the VI [22, 23]. Thus, WOX5, by moving to the VI cells and shootwards, is able to induce *BRAVO* expression in those cells. Additionally, we considered that BRAVO and WOX5 are able to physically interact at the protein level, presumably by binding together, as suggested by Co-IP and FRET-FLIM analysis [28].

While the mobility of WOX5 (possibly through plasmodesmata) has been experimentally tested *in planta* [22, 23], no evidence of intercellular BRAVO transport has been reported. Due to its larger size (BRAVO has a molecular weight of about ~ 36 kDa compared to the ~ 20 kDa of WOX5 [30]), BRAVO proteins are expected to be less mobile than WOX5 proteins, if mobile at all. Moreover, the BRAVO-WOX5 complex, owing to its even larger size, is not expected to move very much from cell to cell. In this respect, while bounds on the plasmodesmata size exclusion limit (SEL) vary, estimates place the SEL lower bound to 27 kDa and upper bound to < 54 kDa for QC/cortex and QC/columella stem cells, being the highest SEL at ~ 60 kDa, between the endodermis/pericycle, pericycle/inner vasculature, and cortical/epidermal cells [31], although these bounds may change depending on different environmental conditions and developmental stages. These values suggest that the BRAVO-WOX5 complex cannot move from cell to cell in the SCN, while WOX5 can.

Taken together, these observations let us propose the following mechanism for BRAVO to confine its own promoter expression (Figure 2A). WOX5 proteins, produced in QC cells and mobile towards the VI cells, are able to activate *BRAVO* expression in the VI. In turn, BRAVO proteins sequester WOX5 into an immobile (from cell to cell) and inactive complex, thus disrupting WOX5 movement and the subsequent activation of *BRAVO* expression there. Hence, the activation of the *BRAVO* promoter by WOX5 is spatially confined by BRAVO proteins, a restriction which becomes released when BRAVO no longer immobilizes WOX5, e.g. in the *bravo* mutant.

**Figure 2:**
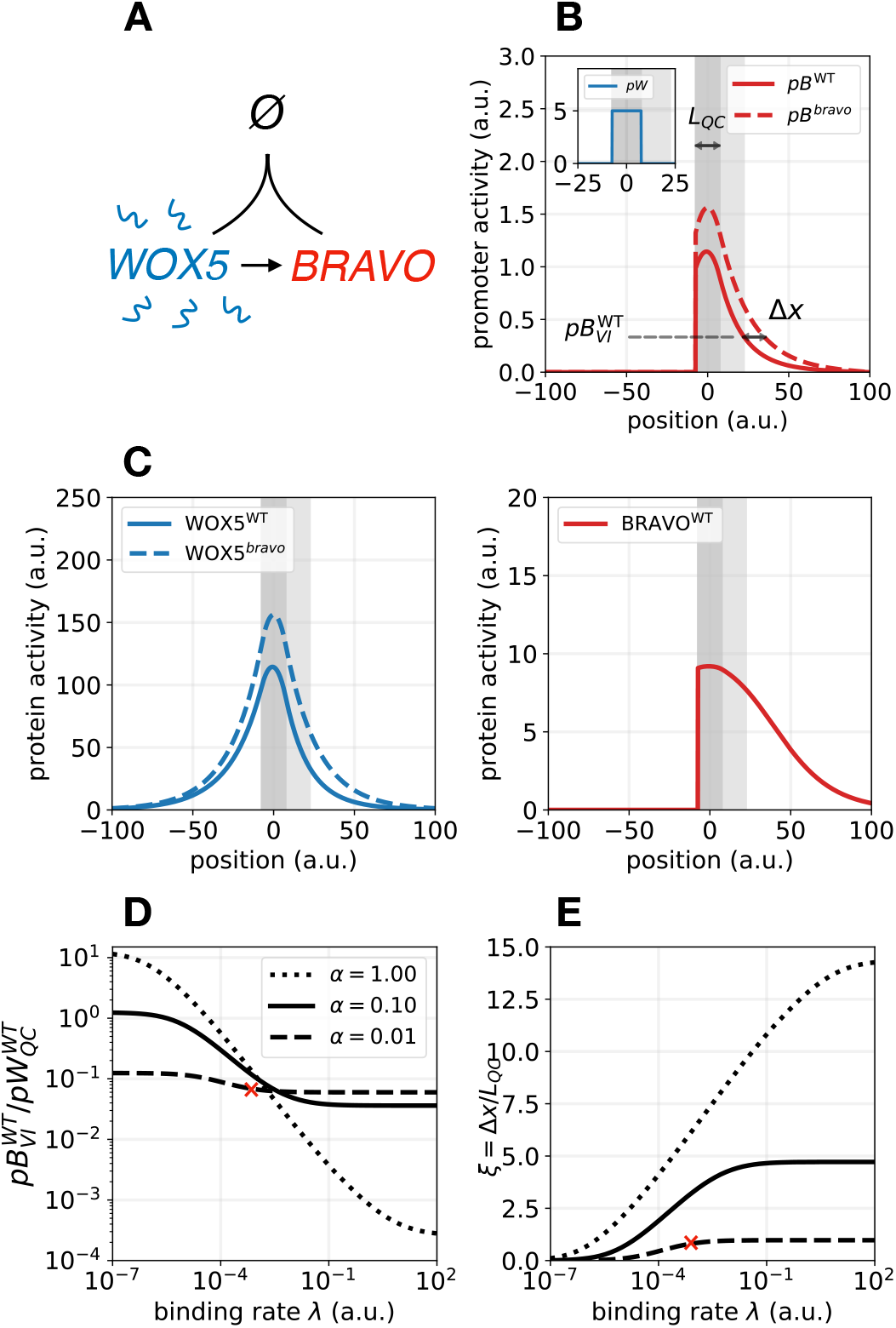
Immobilization by sequestration model. **A)** Cartoon displaying the models’ interactions: WOX5 diffuses (wavy lines) and activates (arrow) BRAVO. In turn, BRAVO immobilizes WOX5 by sequestering it into an immobile and inactive complex (depicted as 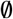). **B)** Stationary profiles of BRAVO and WOX5 promoter activities (*pB* and pW) obtained with this model in the WT (continuous lines) and in the *bravo* mutant (dashed lines). The QC region (dark gray) has a size *LQC* and is where *pW* is active. For simplicity, the VI region (light gray) is defined with this same size. *pB* value in the WT at the end of the VI region is denoted by 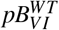. The quantity Δx is a measure of *pB* expansion in the *bravo* mutant as depicted (see Methods for details). If BRAVO confines its own expression in the WT, then Δx >0. **C)** Stationary protein activity profiles of BRAVO (B) and WOX5 (W) in WT (continuous lines) and in the *bravo* mutant (dashed lines) corresponding to the simulations in B). These profiles only depict the proteins not bound to each other. **D, E)** Effect of the binding rate λ and of the BRAVO synthesis rate αon 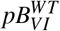 (D) and on Δx (E). In D), the ratio between 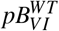 and the constant value of *pW* 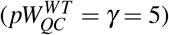 is shown. To be in agreement with experiments [28], this ratio needs to be of order 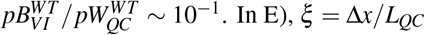. In E), *ξ* = Δx/*L_QC_* represents the expansion of *pB* relative to the QC size. The curves drawn in E) correspond to the same α values as in D). The red crosses in D) and E) mark the values of α and λ used for the simulations in B) and C). Larger values of α increase the expansion but lead to *pB* ≈ *pW* at the QC in the WT (Supplementary Figure 1), which is not experimentally supported [28, 32]. In all panels, parameter values are γ = 5, *d_W_* = *d_B_* = 0.01 and *D_W_* = 4 in arbitrary units (a.u., see Methods and Supplementary Table 1 for further information on parameter values choice). In (B,C) α = 0.01 and λ = 0.001 a.u. Supplementary Figures 1,2 provide supplemental information to this figure.

To evaluate how this mechanism can generate confinement of *BRAVO* expression, we constructed a minimal mathematical model, hereafter named *immobilization by sequestration* model, and studied it in one spatial dimension (Methods, Figure 2A). In this model WOX5 is able to diffuse, while BRAVO is not. Moreover, BRAVO and WOX5 proteins form an immobile complex, i.e. it cannot diffuse. To be consistent with the regulatory interactions between *BRAVO* and *WOX5* reported previously [28], we further require the complex to be inactive, i.e. it does not transcriptionally regulate *BRAVO* nor *WOX5*. Thus, in the model, the production of BRAVO proteins is induced by WOX5, but not by WOX5 when bound to BRAVO. For the sake of simplicity, this activation is assumed to be proportional to the amount of WOX5 proteins. WOX5 is produced at a localized region (dark grey shaded area in Figure 2B) that we identify as the QC, and can diffuse in both directions, namely towards regions that could be identified as the CSC and columella cells (negative values of the position in Figure 2B), or towards the vasculature (positive values of the position in Figure 2B, where the light gray shaded region is identified as VI cells). Conversely, BRAVO can only be activated by WOX5 from the QC and towards the vasculature, but not towards the columella, thus mimicking the asymmetric activity of the BRAVO promoter in the *Arabidopsis* primary root.

In Figure 2B,C we show the stationary activity profiles of BRAVO and WOX5 promoters (pB, *pW*, respectively) and the BRAVO and WOX5 protein concentrations (*B* and *W*, respectively), obtained by numerically solving the model equations in one spatial dimension (Methods). The concentration of the protein complex (which is proportional to the product *BW*) is not shown. The stationary profiles obtained when modeling the *bravo* mutant condition (Methods) are also depicted.

Our results support that the *immobilization by sequestration* mechanism constitutes a plausible way for BRAVO to confine its own expression in the WT (Figure 2B,C). This mechanism still holds when the activation of *BRAVO* expression by WOX5 is not direct but instead occurs through a WOX5 target, as long as this target does not diffuse (Methods).

In this mechanism, binding between WOX5 and BRAVO (mediated by the binding strength *λ*) is necessary for the confinement to take place (Figure 2E). However, high sequestration of WOX5 by BRAVO (large λ and high BRAVO synthesis rate *α*) can lead to an overly exaggerated confinement, with BRAVO being highly expressed at the QC but not in VI cells (Figure 2D, Supplementary Figure 1A), a situation which would be in disagreement with the experimentally observed WT expression (Figure 1B). Hence, the model is able to drive a self-confined expression consistent with that in real roots for low sequestration of WOX5 by BRAVO. Indeed, low sequestration of WOX5 by BRAVO at the QC is expected in the *Arabidopsis* root since the expression of *BRAVO* in the SCN and the amount of BRAVO RNA transcripts are low compared to those of *WOX5* (we take as approximate value 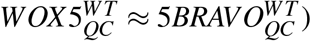 [27, 28, 32].

This mechanism further requires WOX5 to be mobile (Supplementary Figure 1D). The results show that if WOX5 is set to have a large diffusion coefficient, the *BRAVO* expression in the WT becomes larger and much fainter (Supplementary Figure 1C,D), in disagreement with WT expression in Arabidopsis roots [28]. Indeed, WOX5 diffusion has been reported to be rather small [22, 23]. Overall, our results indicate that the *immobilization by sequestration* mechanism is a plausible candidate to explain the observed self-confinement of *BRAVO* expression.

Since sequestration is a necessary ingredient for this mechanism to work, one could argue that the presence of other molecules also binding to BRAVO and/or WOX5 might impair or even destroy the confinement. Indeed, it is known that WOX5 and BRAVO proteins can bind to additional molecules, such as TOPLESS and BES1 [27, 28]. Since these are relatively large molecules (~ 39 kDa and ~ 124 kDa, respectively [30]) it would be possible for them to immobilize WOX5. Therefore, these other molecules can also generate confinement of *BRAVO* expression, as confirmed by modeling this scenario (Supplementary Figure 2). If these molecules sequester BRAVO or WOX5 excessively, preventing the binding between the two, the mechanism of BRAVO self-confinement becomes compromised (Supplementary Figure 2). Therefore, in order to maintain the self-confinement of *BRAVO* expression in the presence of these other factors, their sequestering effect on BRAVO and WOX5 must be small.

Altogether, the *immobilization by sequestration* mechanism constitutes a plausible candidate for explaining the self-confined *BRAVO* expression to both the QC and VI cells in the WT. Notably, the WT stationary profile of WOX5 proteins (W, not bound to BRAVO) obtained for this model is not sym-metric, but decays differently above and below the QC (Figure 2C). This behaviour is caused by the presence and absence, respectively, of BRAVO proteins in the two distinct spatial regions. Above the QC, the gradient of WOX5 proteins is steeper than below the QC, where BRAVO cannot be activated. Hence, the presence of BRAVO makes the WOX5 gradient more abrupt, restricting the spatial domain where WOX5 proteins are concentrated, and consequently confining the *BRAVO* expression domain (recall that in this simplified model *BRAVO* expression is proportional to WOX5 levels). Therefore, this mechanism not only drives self-confined *BRAVO* expression but also results in a confined action of WOX5 proteins.

### 2.2 BRAVO cannot confine its own expression domain in the root SCN only by inactivating WOX5

The self-confinement of *BRAVO* expression in the *immobilization by sequestration* model requires WOX5 to diffuse. Experiments suggest that, while mobile, WOX5 does not move very large dis-tances [22, 23]. Hence, its small diffusion may not be sufficient to explain the self-confinement of *BRAVO* expression observed in real roots. Since activation of *BRAVO* by WOX5 could happen through intermediary molecules, we asked whether the binding between the two proteins could still be sufficient to drive self-confinement of *BRAVO* expression if WOX5 is activating *BRAVO* not directly, but through a highly mobile WOX5 target, hereafter named *X*. This mechanism also assumes that the BRAVO-WOX5 complex is transcriptionally inactive, and hence BRAVO, by binding to WOX5, prevents WOX5 from activating *X* (Figure 3A). To evaluate this scenario, we again formulated a minimal model, hereafter named *attenuation by sequestration* model (Methods), and studied its implications by numerically simulating WT and *bravo* mutant backgrounds, as done in the previous section. In this minimal model, neither WOX5, nor BRAVO nor the complex can diffuse.

**Figure 3:**
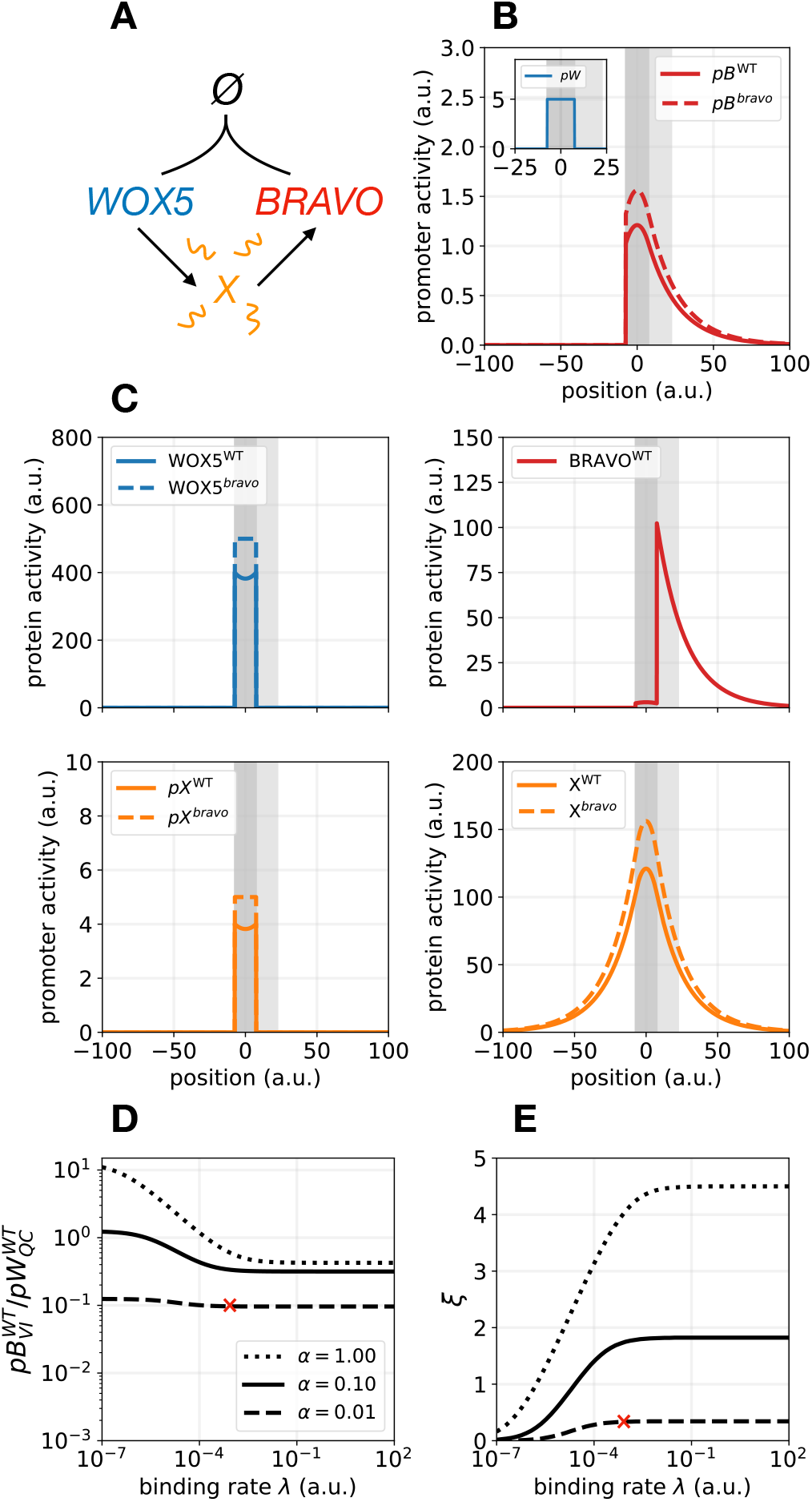
Attenuation by sequestration model. **A)** Cartoon of the interactions: WOX5 activates a highly diffusible intermediary *X*, which in turn activates BRAVO. BRAVO sequesters and inactivates WOX5, preventing the activation of *X*. **B)** Stationary activity profiles of *pB* and *pW* in WT (continuous lines) and in the *bravo* mutant (dashed lines). **C)** Stationary profiles of BRAVO (B), WOX5 (W) and *X* protein activities, as well as promoter activity *pX*, in WT (continuous lines) and in the *bravo* mutant (dashed lines) corresponding to the simulations in B). **D, E)** Effect of the binding rate *λ* and of the BRAVO synthesis rate *α* on 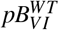 (D) and on Δx (E). In D), the ratio between 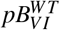 and the constant value of *pW* is shown 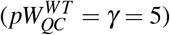. In E) *ξ* = Δx/*L_QC_* is shown with same *α* values as in D). Red crosses mark the values of *α* and *λ* used for the simulations in B) and C). In all panels *γ* = 5, *β* = 0.01, *d_W_* = *d_B_* = *d_X_* = 0.01 and *D_X_*= 4 in arbitrary units (a.u., Methods, Supplementary Table 1). In (B,C) *α* = 0.01 and *λ* = 0.001 a.u. Supplementary Figure 3 provides supplemental information to this figure.

The results confirmed that the *attenuation by sequestration* mechanism is also able to drive BRAVO self-confinement (Figure 3B). In this mechanism, because WOX5 is sequestered by BRAVO, *X* becomes less activated at the QC and hence reaches with high concentration smaller spatial regions, compared to the case when BRAVO is absent (Figure 3B). However, the results show that this mechanism requires WOX5 to be highly sequestered by BRAVO to drive a noticeable self-confinement (Figure 3B and Supplementary Figure 3A-D). This implies that *BRAVO* has to be strongly expressed (similarly to *WOX5*) at the QC (Supplementary Figure 3). In *Arabidopsis* roots, since *WOX5* is much more strongly expressed than *BRAVO* [28] (Figure 1B,C), we expect a low sequestration of WOX5 by BRAVO. This suggests that the *attenuation by sequestration* mechanism is not relevant to account for *BRAVO* self-confined expression in the *Arabidopsis* primary root SCN.

### 2.3 BRAVO can confine its own expression domain by repressing its mobile activator

The *attenuation by sequestration* mechanism indicates that a transcription factor can confine its own expression by reducing the production of its mobile activator. Based on this, we envisaged a third scenario which does not have the limitations imposed by sequestration. In this case, BRAVO represses the production of its activator, *Z*, which is mobile and is activated by WOX5 (Figure 4A). We call this the *repression* model (Methods). Reported transcriptomics of QC cells have revealed that most of the genes whose mRNA levels are de-regulated in *wox5-1* and *bravo-2* mutants show opposite regulations [28], thus opening the possibility of one of these genes to act as the intermediate factor *Z*, which in our model is downregulated in the *wox5* mutant but upregulated in the *bravo* mutant. Hence, transcriptomic data in the QC [28] suggest several candidates for *Z*.

**Figure 4:**
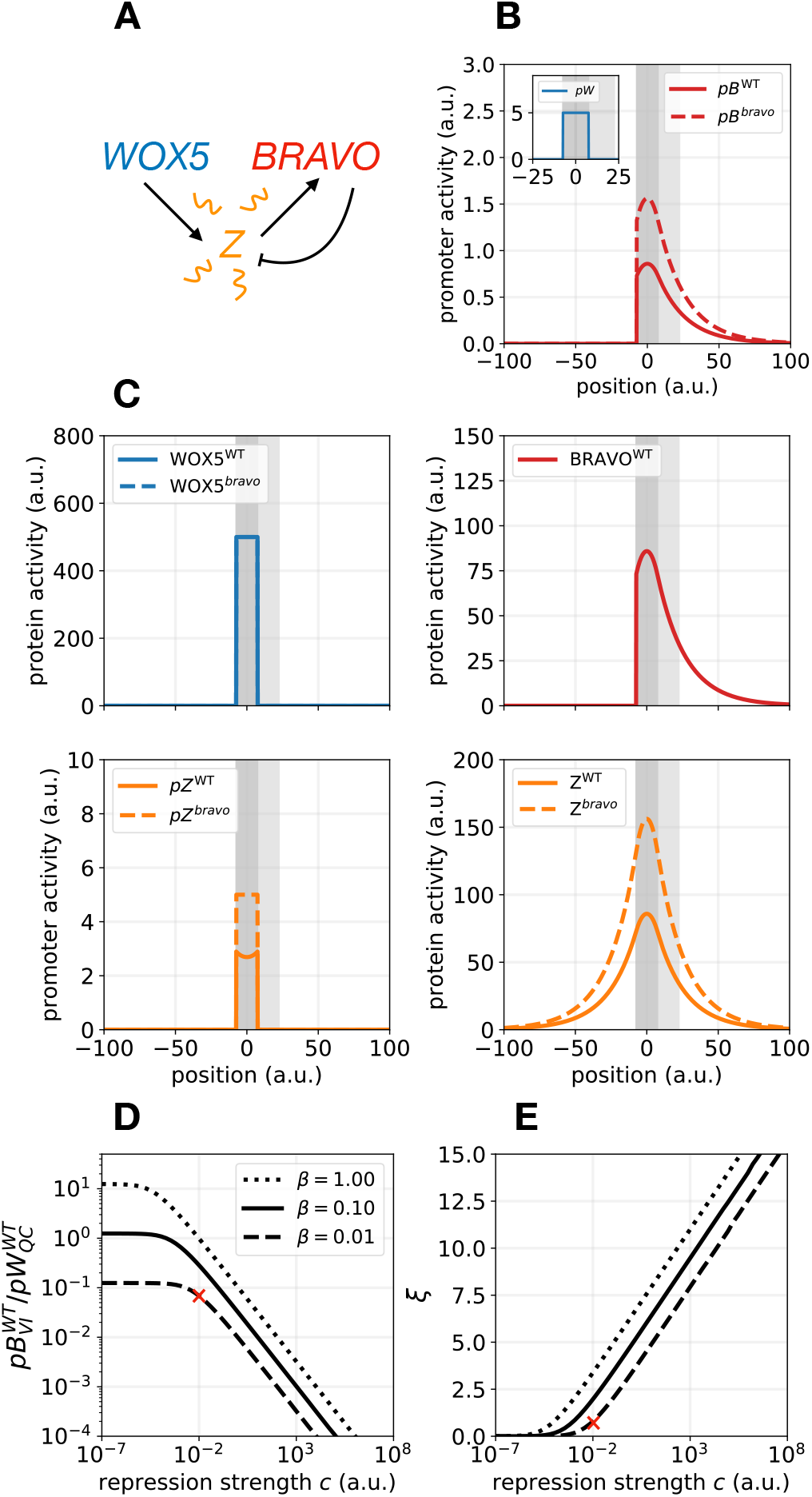
Repression model. **A)** Sketch of the interactions: WOX5 activates a diffusible factor Z, which activates BRAVO. BRAVO, in turn, is able to repress (blunt arrow) the activity of Z. **B)** Stationary activity profiles of *pB* and *pW* in WT (continuous lines) and in the *bravo* mutant (dashed lines) obtained with the repression model. **C)** Stationary profiles of BRAVO (B), WOX5 (W) and *Z* protein activities as well as promoter activity *pZ*, in WT (continuous lines) and in the *bravo* mutant (dashed lines) corresponding to the simulations in B). **D, E)** Effect of the repression strength *c* and of the *Z* synthesis rate *β* on 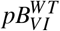 (D) and on Δx (E). In D), the ratio between 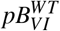 and the constant value of *pW* is shown 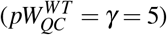. In E) *ξ* = Δx/*L_QC_* is shown with same *β* values as in D). Red crosses mark the values of *c* and *β* used for the simulations in B) and C). Larger *β* and *c* values drive larger expansions but involve a very strong increase of *pB* levels in the *bravo* mutant (see Supplementary Figure 4), which is not in agreement with experiments (Figure 1). In all panels parameter values are *α* = 0.01, *γ* = 5, *d_W_* = *d_B_* = *d_Z_*= 0.01 and *D_Z_* = 4 in arbitrary units (a.u., Methods, Supplementary Table 1). In (B,C) *β* = 0.01 and *c* = 0.01 a.u. Supplementary Figure 4 provides supplemental information to this figure.

Simulations indicate that the *repression* mechanism is also able to induce the confinement of *BRAVO* expression (Figure 4B) and, as with previous mechanisms, shows that the action of WOX5, mediated by its target Z, becomes spatially restricted in the WT, while it expands in the *bravo* mutant (Figure 4B,C).

The *repression* mechanism relies on the repression of *Z* by BRAVO (Figure 4E) and on the mobility of *Z* (Supplementary Figure 4). Albeit movement of *Z* is required for this mechanism to drive self-confinement, simulations indicate that large diffusion coefficients of *Z* drive a *BRAVO* expression in the WT that is too faint and too spread to be compatible with the GFP expression in *Arabidopsis* root SCN (Supplementary Figure 4, Figure 1). This suggests that the diffusion coefficient of *Z* should be small.

As opposed to the *attenuation by sequestration* mechanism, the *repression* mechanism does not require a very strong *BRAVO* expression in the WT to drive noticeable self-confinement (i.e. the concentration threshold of BRAVO to repress *Z* is low). Accordingly, *BRAVO* expression values lower than those of *WOX5*, as seen in the Arabidopsis root SCN, are consistent with strong self-confinement through repression (Figure 4B). Further analysis of the model shows that strong repressions enhance the self-confinement, but can result in unrealistic *BRAVO* expression profiles in the WT, limited only to the QC and not reaching the VI cells (Figure 4D,E). In addition, strong repression involves not only spatial confinement but also a dramatic reduction of *BRAVO* expression at the QC in the WT scenario compared to the *bravo* mutant (Supplementary Figure 4), a situation which is not observed experimentally [28](Figure 1B,C). Taken together these results indicate that if the *repression* mechanism takes place at the SCN of *Arabidopsis* to confine the *BRAVO* expression domain, then BRAVO represses the WOX5 target *Z* only weakly.

### 2.4 The immobilization by sequestration mechanism is sufficient and the re-pression mechanism enhances self-confinement in *Arabidopsis*

Our previous results suggest that both the *immobilization by sequestration* and the *repression* mechanisms are conceivable candidates to explain the self-confined expression of *BRAVO* in the *Arabidopsis* root SCN. However, it remains unclear whether the diffusion of WOX5 is sufficiently large to make the *immobilization by sequestration* mechanism sufficient, and whether the weak repression required in the *repression* model is enough to produce the confinement observed in real roots. To address these issues, we modelled these mechanisms on realistic root layouts, where the cellular geometry of the roots is explicitly incorporated. We assumed both mechanisms to be present at the same time (which incidentally also incorporates the *attenuation by sequestration* mechanism as a side-effect) (Figure 5A, hereafter named *mixed model*). WOX5 can move between cells and activates a mobile target *Z*, which activates BRAVO, and BRAVO feeds back on *Z* to repress its activity. BRAVO and WOX5 are able to bind together, forming a transcriptionally inactive complex which is not able to activate *Z* and which, like BRAVO, cannot move from cell to cell. We next evaluated this mixed model in realistic root geometries to assess whether it can predict the changes in expression observed in *Arabidopsis* roots. Besides modeling the dynamics of BRAVO and WOX5, we also modelled the dynamics of GFP proteins produced by the BRAVO and WOX5 promoters, denoted by *pBRAVO:GFP* and *pWOX5:GFP*. For GFP molecules, the production was set to be proportional to their corresponding promoter activities, and their degradation was set to be linear (just like the others). In addition, GFP molecules were assumed to be mobile, both within and between cells.

**Figure 5:**
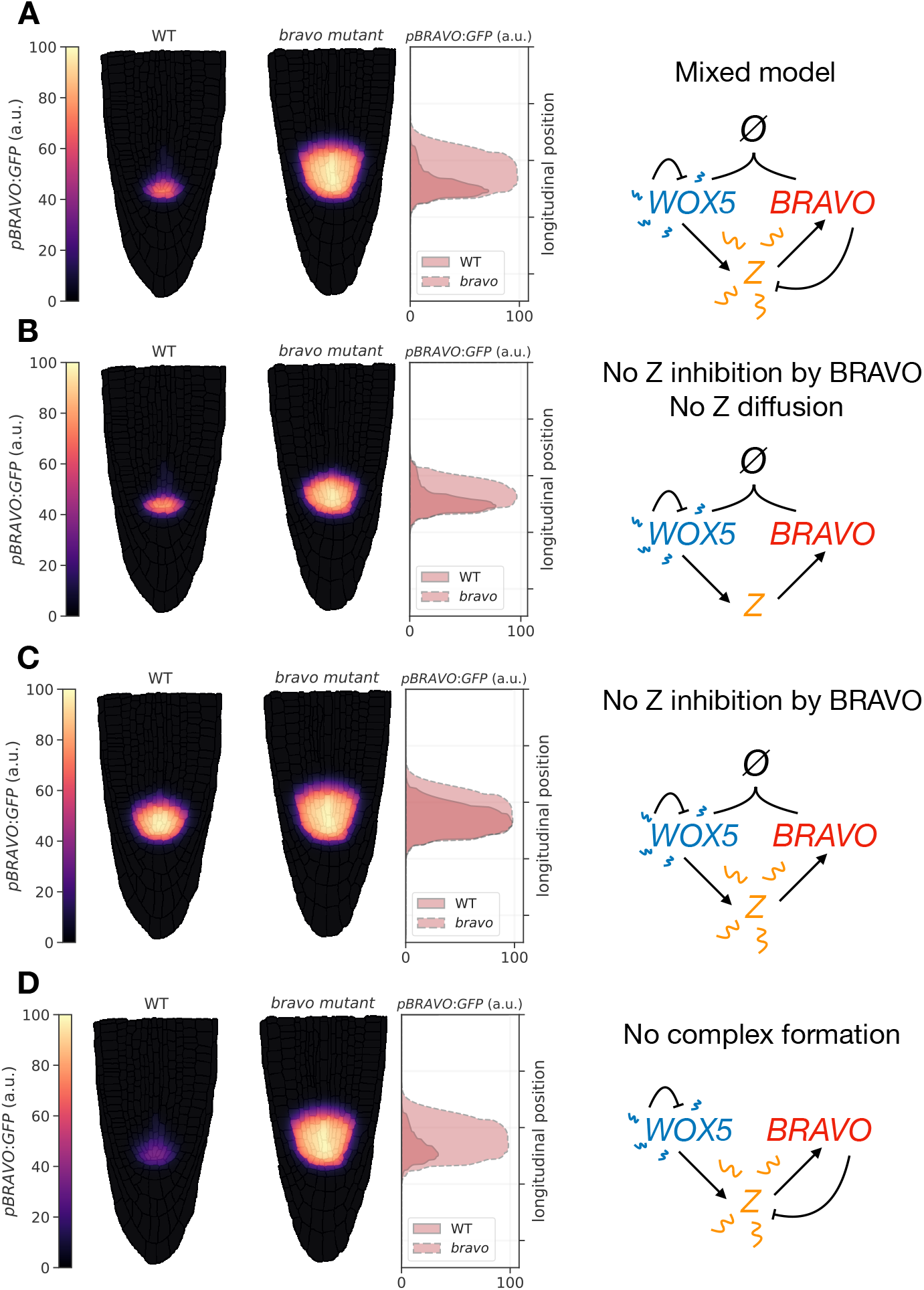
*pBRAVO:GFP* simulated in realistic root layouts. Simulation results for the stationary activity of *pBRAVO:GFP* in WT (left root) and in *bravo* mutant (right root) for different regulatory interactions (each depicted as a cartoon on the right). The middle panel depicts *pBRAVO:GFP* along the midline longitudinal (vertical axis). **A)** Mixed model: WOX5 is able to diffuse, self-repress and activate a highly mobile intermediary Z. This intermediary activates BRAVO, which in turn feeds back on *Z* to repress its activity. BRAVO and WOX5 can bind into an immobile and inactive complex. In the WT, *pBRAVO:GFP* is confined to the SCN, and it is expanded in the *bravo* mutant. The stationary profiles of all other molecules are depicted in Supplementary Figure 9. **B)** Confined expression of *pBRAVO:GFP* can still be obtained when *Z* does not diffuse and is not repressed by BRAVO (this corresponds to the *immobilization by sequestration* mechanism). The expansion in the *bravo* mutant is smaller than for the mixed model. **C)** The mixed model without the repression of *Z* by BRAVO makes *pBRAVO:GFP* to be more expanded in the WT, and with similar absolute levels as in the *bravo* mutant. Hence, little confinement is achieved in the WT. **D)** If there is no binding between BRAVO and WOX5, the activity of *pBRAVO:GFP* in the WT dramatically decreases due to stronger repression through *Z*. In all panels, cell walls are superimposed (in black with transparency) over the colormap so they can be easily visualized. Parameter values in Supplementary Table 3. Supplementary Figures 5–9 provide supplemental information to this figure.

The implementation of the mixed model in two-dimensional root geometries considered the realistic shape of cells and the presence of cell walls. The geometries of WT and *bravo* mutant roots were considered separately, by using different layouts. The main features included in this new framework can be enumerated as follows (for further details, see Methods and SI).

1. The spatial discretization of the realistic root layout was made at the pixel scale. Hence, the size of the cytoplasm and cell walls is determined by the number and localization of their corresponding pixels (Supplementary Figure 5).
2. The dynamics of the molecular components in the interior of the cells is distinct to the dynamics in the cell walls. Specifically, in all the pixels belonging to the cell’s interior, molecules can be produced, regulated, degraded, can sequester other molecules and are able to diffuse. Inside the cell walls, however, molecules are only able to diffuse.
3. The diffusion coefficient in the cell wall is set to be smaller than in the interior of the cells. With this assumption the physical boundaries between cells are naturally incorporated, leading to discontinuities in the concentration profiles of molecular factors. These are to be expected for any molecule diffusing freely inside the cytoplasm but moving only occasionally from cell to cell, as it may happen, for example, in plasmodesmata-mediated transport. For simplicity (and lack of evidence), we assume that BRAVO and the BRAVO-WOX5 complex can only move inside the cell, and are unable to diffuse through cell walls. Conversely, WOX5 and *Z* proteins can move inside the cytoplasm with diffusion coefficients 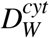 and 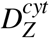, respectively, and between cells with diffusion coefficients 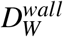 and 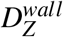, respectively. The diffusion coefficient of GFP and WOX5 proteins was assumed to be similar, both in the cytoplasm and in the cell wall, since both proteins have aminoacid sequences of similar lengths (~ 20 kDa for WOX5 and ~ 27 kDa for GFP [30]). Because GFP proteins are able to diffuse from cell to cell, they can reach cells where there is no promoter activity. However, the diffusion at the cell wall is set to be sufficiently low so that this effect is small.
4. Cells are classified by cell type. We defined nine different cell types: quiescent center, vascular initials, vascular cells, cortex, endodermis, epidermis, columella stem cells, columella cells, and lateral root cap cells (Figure 1A). This classification enables setting different dynamics for the proteins in distinct cell types, as well as different transport properties (through different diffusion coefficients). We set the activation of BRAVO by WOX5 only in the QC, vascular initials, vascular cells, cortex and endodermis, but not in the remaining cell types (Supplementary Figure 6). This last assumption is based on the experimental observation that *BRAVO* is expressed only in inner tissues, from the SCN upwards. To account for the expression of BRAVO promoter in some VI and V cells in the *wox5* mutant [28] (Figure 1D), basal production of BRAVO, independent of WOX5, is set in few of these cells (Supplementary Figure 7). For simplicity, WOX5 is only allowed to be produced at the QC (albeit recent observations show that low values of promoter expression are also present in the vascular initials [22, 23]). Finally, diffusion and degradation can occur in all cell types.

To avoid the possible drawbacks caused by linear synthesis rates used in the minimal models presented (such as the lack of saturation and thresholds of activity), herein we used transcriptional regulations described by Hill functions, with basal activity and saturation values (Methods). In order to account for the increased *WOX5* promoter expression in the *wox5* mutant, we included a negative feedback on WOX5, a regulation that, in turn, induces a decrease in WOX5 activity in the *bravo* mutant scenario, as sequestration of BRAVO by WOX5 allows the WOX5 promoter to be less repressed in the WT [28].

Assuming biologically realistic parameters for diffusion, production and degradation of molecules (see Supplementary Table 3), we found that the mixed model can explain the behaviour of *pBRAVO:GFP* in the WT and the expansion of its domain in the *bravo* mutant (compare Figure 5A with Figure 1B,C). Here, WOX5 diffusion is set to be low enough such that its promoter activity, in the absence of any regulatory factor, is mostly localized at the QC, VI and CSC (Supplementary Figure 8), as found experimentally. The simulations show that BRAVO self-confinement additionally induces WOX5 proteins and the diffusible target *Z* to be confined (Supplementary Figure 9).

The small diffusion of WOX5 is sufficient to drive a realistic confinement of *pBRAVO:GFP* through the *immobilization by sequestration* mechanism alone (Figure 5B). However, this only happens if activation of BRAVO by WOX5 occurs through a non-diffusible WOX5-target *Z* (Figure 5C). For a diffusible *Z*, the *repression* mechanism is required to drive BRAVO self-confinement (Figure 5A,C). Under these circumstances, sequestration between BRAVO and WOX5 facilitates a higher *pBRAVO:GFP* expression in the WT (Figure 5A,D).

Taken together, the results in the realistic root layout support that the immobilization of WOX5 by BRAVO is sufficient for the self-confined expression of *BRAVO* in the root SCN.

## 3 Discussion

We have previously shown that BRAVO and WOX5 regulate each other expressions and that their binding into a protein complex can be relevant for these regulations and for BRAVO and WOX5 action on QC divisions [28]. Here we have shown that these interactions, together with the diffusion of WOX5, are sufficient to account for the spatially self-confined expression of *BRAVO*, as revealed by the *immobilization by sequestration* mechanism. Furthermore, the opposite regulation of common targets by WOX5 and BRAVO, as inferred by transcriptional profiling [28], support the complementary scenario where this confinement is induced by a negative feedback between BRAVO and a factor activated by WOX5 (the *repression* mechanism).

These mechanisms establish a negative feedback of BRAVO on itself. As such, they mechanistically account, at least partially, for the effective negative self-regulation of BRAVO expression in the whole SCN previously proposed [28]. Moreover, the results herein indicate that this negative regulation is dependent on WOX5.

To obtain these results we investigated three regulatory mechanisms for self-confined expression (*immobilization by sequestration*, *attenuation by sequestration* and *repression*). These three mechanisms have in common that the emergence of self-confinement involves a negative feedback with a mobile activator and an immobile inhibitor. Each of the three mechanisms can be understood as a different regulatory way for this negative feedback to be accomplished: by immobilizing or by reducing, through sequestration or repression, the production of the activator. Therefore they can be placed within a general framework of self-confinement, which could go under the generic name of *attenuation of a mobile activator*. All these mechanisms drive the self-confinement of both the repressor and the activator. Yet, the three mechanisms are not equivalent, each of them having their distinct characteristics, as our analysis revealed. The most noticeable feature is perhaps the fact that the immobilization mechanism involves a change in the gradient profile of the activator (Figure 2D, SI Text) whereas the other mechanisms do not.

A different mechanism to drive self-confined expression has been proposed for WUSCHEL in the shoot apical meristem of *Arabidopsis*. According to it, WUSCHEL protein confines its own expression by activating its repressor, CLAVATA3, which is highly mobile [12, 33]. Both this mechanism and the *attenuation of a mobile activator* studied here have in common that a negative feedback is responsible for the self-confinement. However, in the *attenuation of a mobile activator* the strongly mobile component is the activator and not the repressor. The generic process of *attenuation of a mobile activator* is thus a distinct mechanism for self-confinement and, because of its minimal assumptions, we expect it, or a variant thereof, to take place in very distinct developmental contexts. For instance, Hedgehog signaling in the Drosophila wing confines its own expression by activating one of its repressors, the nuclear zinc finger *Master of thickveins* [34]. Thus, this Hedgehog self-confinement may be framed within the *attenuation of a mobile activator* mechanism, where Hedgehog signaling acts as the mobile activator (*Z* in the repressor model) and *Master of thickveins* acts as the immobile repressor.

The mechanism of *immobilization by sequestration* can be related to mechanisms for robust morphogen gradient profiles [35, 36]. Morphogens are ligand molecules that are produced at localized sources but can diffuse, generating an activity gradient that can then be interpreted by different genes, activating or repressing them in a concentration-dependent manner. It has been proposed that the sequestering of the morphogen by receptors might lead to an effective non-linear degradation of the ligand, resulting in concentration profiles that are robust to changes in the rate of production at the source [35]. We could make the correspondence between such models and ours by identifying WOX5 as the morphogen and BRAVO as the receptor. In agreement with what has been described for morphogen gradients induced by non-linear degradation [35], we find that the gradient of WOX5 decays much more abruptly in the presence of BRAVO than without, thus suggesting that the specific regulations between BRAVO and WOX5 may be tuned to achieve robust activity profiles.

Modeling of root tissues has been most commonly done in terms of simplified rectangular geometries in which cells and cell walls are subdivided in squares or rectangles [37, 38], or by considering cellular layouts with diffusing molecules between but not within cells [39, 12, 40]. By using pixels as the basic unit for discretization, we are able to model the shapes of cells in a realistic manner, and consider both the interactions within and between cells. A similar pixel-based approach has been used to model hormonal crosstalk in the *Arabidopsis* root [41]. Mathematically, our framework can be characterized as a reaction-diffusion model in heterogeneous media, where the spatial inhomogeneities appear due to the presence of cell walls, which involve different diffusion coefficients. The realistic root layout used for the simulations can be extended to include internal structures within the cells (such as the nucleus) as well as structures in the cell walls (e.g. specific communication channels). Therefore, it has the potential to implement and evaluate much more complex scenarios in a manageable way.

The similarities between stem cell niche organization in animals and plants may represent the outcome of convergent evolution [42]. Multicellularity – a necessary condition for stem cell niches to emerge – is thought to have evolved independently in both kingdoms [43, 44], implying that their presence in widely disparate organisms may be a direct consequence of developmental constraints and not of historical contingencies. Similar mechanisms of niche regulation are therefore to be expected, not through common genes or molecules, but through more general regulatory principles. The fact that stem cell niches consist on narrow regions of few cells clustered together, in opposition to large numbers of cells distributed over the whole organism, possibly emerged as a way to ensure a proper balance between centralized renewal and genome integrity, by minimizing deleterious mutations which may be able to spread across whole cell lineages [17, 45]. Indeed, the smaller the population of stem cells and the lower their division rate, the less likely for deleterious mutations to accumulate in differentiated tissues [46]. The mechanism proposed in this paper establishes a balance between the communication mediated by the activator (WOX5) with confining its action, ensuring communication remains local.

In the Arabidopsis SCN, these signals allow cells to communicate between them and with other cell types, at the same time as they create boundaries within which local information can be transmitted. Indeed, QC cells have been shown to influence neighboring cell types such as the CSC, where WOX5 can play a crucial role as a signaling agent [21]. We propose that towards the vasculature, *BRAVO* can be a signaling molecule, which, by actively restraining its own expression from reaching cells far away from its source, ensures that the small microenvironment of the SCN remains confined within the root. The molecular processes underlying this spatial restriction and their implications for proper stem cell renewal are just beginning to be uncovered. Mechanisms like the ones proposed here involve very general principles which contribute to the understanding of stem cell populations not only in plants, but in multicellular organisms on the whole.

## 4 Methods

The models formulated set the rate of change of protein concentrations of BRAVO (*B*) and WOX5 (*W*), by using partial differential equations where the transport of the mobile proteins is modelled through diffusion. In the models where intermediate factors are present (*X* in the attenuation by sequestration model and *Z* in the repression model), their dynamics is also considered. We only focus on the stationary solutions of the models, assuming these account for the experimentally reported expressions.

In all the models, the rate of synthesis of each protein is assumed to be proportional to its corresponding promoter activity and the quasi-stationary approximation for mRNAs (i.e. mRNAs dynamics are assumed to be very fast compared to the dynamics of proteins) is done (see SI Text). For simplicity, proteins are assumed to degrade linearly. The rate at which two proteins form a complex is assumed to be proportional to the product between the two protein concentration variables (e.g. ∝ *BW* for the BRAVO-WOX5 complex). Therefore, we only consider pair-wise interactions, omitting higher order reactions. The complex is assumed to bind reversible and to degrade linearly, with very fast dynamics enabling its quasi-steady state approximation. As a result, the concentration of complex is not explicitly computed, but only the amounts of not bound proteins. Since all results are computed at the stationary state, this does not introduce any additional approximation. Finally, complexes are taken to be unable to transcriptionally regulate any of the proteins considered (for simplicity we name them inactive complexes, albeit they could regulate other factors not modelled herein). SI Text contains further details on the derivation of the models equations from the full set of equations which include mRNAs and complexes. This modeling approach is analogous to [28].

Subsections 4.1, 4.2 and 4.3 describe the one-dimensional minimal models in WT scenarios, while the dynamics of *bravo* mutants are described in section 4.4. Section 4.5 describes how the BRAVO stationary profiles obtained from the minimal models in the WT and the *bravo* mutant are compared, as well as the constraints imposed by experimental data on the parameter values. Section 4.6 details the construction of the realistic root layout. Section 4.7 describes the equations used for the simulations of the mixed model in the realistic root layout. Section 4.8 explains the numerical details of all models simulated.

### 4.1 Immobilization by sequestration mathematical model

In this case, WOX5 activates BRAVO, while both proteins are able to form an inactive complex, which is rapidly degraded. For simplicity, the activation of BRAVO by WOX5 is set to be linear. In the WT, the dynamics of *B* and *W* are:

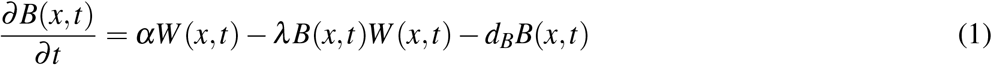

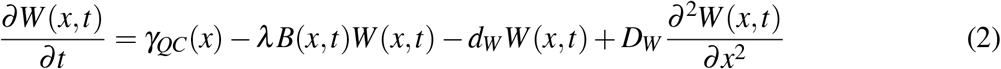

where *x* denotes the spatial position in one dimension and *t* denotes time. Here *B* and *W* stand for the BRAVO and the WOX5 protein concentrations, respectively, when are not bound to each other, and the complex they form is not explicitly modelled as a variable (see SI Text). The parameter *α* measures the production rate of BRAVO per unit concentration of WOX5, and due to the linearity of the promoter, has units of inverse time. WOX5 is produced at a constant rate *γ*_*QC*_(*x*), where the subscript and the explicit spatial dependence indicate that it is only produced at the QC. We choose *γ*_*QC*_(*x*) to be a rectangular function of the form

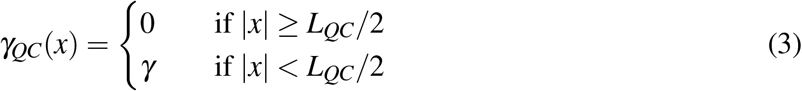

where *L_QC_* is the total length of the QC region. This implies that WOX5 production only occurs in the region delimited by 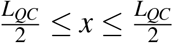, with constant production rate *γ*. Degradation of BRAVO and WOX5 is controlled by the parameters *d_B_* and *d_W_*, respectively. Complex formation between BRAVO and WOX5 is mediated by the parameter *λ*, which sets the rate at which the two factors interact per concentration unit of each of them and involves binding, unbinding and degradation rates (SI Text). Finally, WOX5 is able to diffuse from the QC with rate *D_W_*. As explained in detail in SI, in order to have confinement through the immobilization by sequestration mechanism, it is essential for the formation of the BRAVO-WOX5 complex to be either irreversible, or reversible but being subject to degradation. The *immobilization by sequestration* model constitutes a simplified but spatially dependent version of the *complex formation model* proposed in [28] to explain the regulations between BRAVO and WOX5 in the whole *Arabidopsis* stem cell niche.

The promoter activities of BRAVO and WOX5, which are computed at the stationary state (i.e. when all time derivatives are zero), are defined as *pB*(*x*) = *αW*_*s*_(*x*) and *pW*(*x*) = *γ*_*QC*_(*x*), respectively, where *W_s_*(*x*) denotes the spatial profile of WOX5 in the stationary state, as indicated by the subscript s. We also refer to these activities as promoter expressions.

Another version of this model, where an additional sequestrator affects the dynamics of BRAVO and WOX5, is described in SI (with results in Supplementary Figure 2).

### 4.2 Attenuation by sequestration mathematical model

In this second scenario WOX5 activates an intermediary factor *X*, which in turn activates BRAVO. *X* is able to diffuse whereas WOX5 is not. BRAVO and WOX5 form a complex, assumed to be inactive and immobile. The dynamics of *B*, *X* and *W* in the WT are:

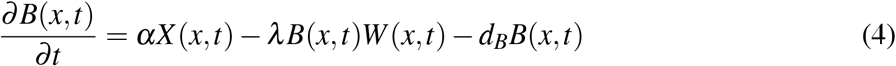

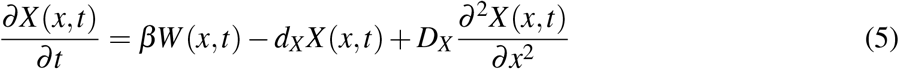

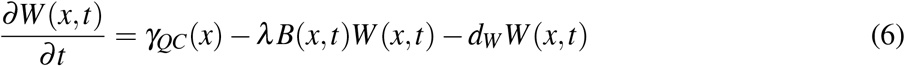

As in the previous model, *B* and *W* stand for the proteins not bound to each other. The new parameters *β*, *d_X_* and *D_X_* characterize the production, degradation and diffusion of the intermediary factor *X*, respectively. In this model, the BRAVO promoter in the stationary state is denoted by *pB*(*x*) = *αX*_*s*_(*x*), where *X_s_*(*x*) denotes the concentration profile of the intermediary in the stationary state, while the WOX5 promoter remains as in the previous case, *pW*(*x*) = *γ_QC_*(*x*), with *γ_QC_* only affecting the QC region (as in Eq.(3)). SI describes an alternative version of this model where WOX5 is allowed to diffuse (results shown in Supplementary Figure 3).

### 4.3 Repression of a mobile activator mathematical model

In this case, WOX5 activates an intermediary factor *Z*, which activates BRAVO. In turn, BRAVO feeds back on *Z* by repressing it. *Z* diffuses whereas WOX5 and BRAVO do not. Binding between BRAVO and WOX5 is not present. The dynamics of *B*, *Z* and *W* in the WT are:

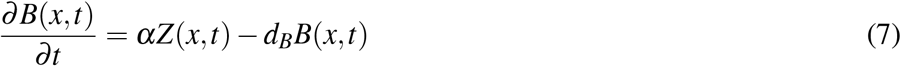

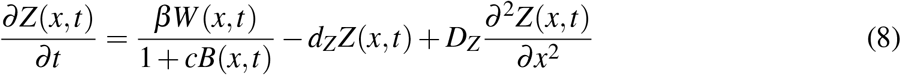

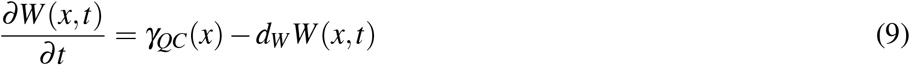

The parameters *β*, *d_Z_* and *D_Z_* describe the production, degradation and diffusion of *Z*, respectively, while the new parameter *c* sets the threshold of *Z* repression by BRAVO. In this case, promoter activities in the stationary state are given by 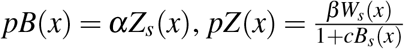 and *pW*(*x*) = *γ_QC_*(*x*) (defined by Eq.(3)), where again the subscript *s* indicates that the concentration profiles are those corresponding to the stationary state.

### 4.4 Modeling *bravo* mutants

The same type of approach as in [28] is used to simulate the *bravo* mutant. Specifically, to model this mutant, the very same dynamical equations and the same parameter values are used as those to model the WT, except for BRAVO which is set as *B*(*x, t*) = 0 for all *x* and *t*. This leads to stationary values of *W_s_*(*x*), *X_s_* (*x*) and *Z_s_*(*x*) that are different than in the WT. While no dynamical equation for *B*(*x, t*) is set, there is a promoter activity of BRAVO in the stationary state, *pB*(*x*), which is as defined for the WT but with the stationary profiles of the mutant. We exemplify this with the immobilization by sequestration model. Setting *B*(*x, t*) = 0 into equations (1,2), we get:

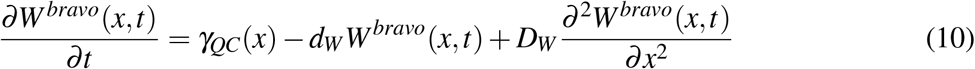

where the new superscript *bravo* indicates that the solution of the equation corresponds to the *bravo* mutant. The stationary BRAVO promoter is 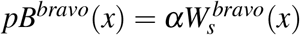. The same procedure is applied for the other models.

### 4.5 Measures of expansion and intensity of simulated *BRAVO* expression

For simplicity, in the one-dimensional minimal models, the VI region (light gray area in Figures 2B,C, 3B,C, 4B,C) is defined to be of the same size as the QC region (*L_QC_*, *x* ∈ [—*L_QC_*/2, *L_QC_*/2]). Hence, the end of the VI is at position *x_VI_* = 3*L_QC_*/2. To quantify whether the simulation results of the minimal models show a stationary BRAVO promoter expression more extended in the *bravo* mutant than in the WT, we compute the value of the stationary BRAVO promoter expression in the WT at the end of the VI region and define this value as 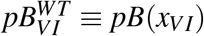. We then compute the spatial position at which the stationary BRAVO promoter expression in the *bravo* mutant takes this value, and define this position as *x^bm^* (i.e. *x^bm^* is defined as 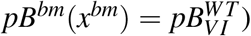. Then, we define Δ*x* ≡ *x^bm^* – *x_VI_* and use it as the measure of how much the BRAVO promoter expression in the *bravo* mutant is expanded (Δx >0) or contracted (Δ*x* < 0) compared to its expression in the WT. This change is then normalized to the size of the QC region, by defining the non-dimensional parameter *ξ*:

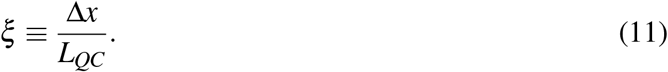

Thus, *ξ* quantifies the change of stationary BRAVO promoter expression in the *bravo* mutant relative to the size of the QC, i.e. *ξ* = 1 means that the stationary BRAVO promoter expression in the *bravo* mutant is expanded a region as large as the QC compared to the WT.

We also computed for the WT the ratio between the stationary BRAVO promoter expression at the end of the VI and the stationary WOX5 promoter expression at the QC (*x* = 0):

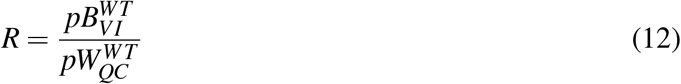

To make the connection with experimental data, we impose that this quantity must remain within a certain interval close to *R* ≃ 0.05 – 0.1, which is within the range observed in experimental data [28, 32]. Moreover, to be in agreement with *BRAVO* and *WOX5* expression data in WT Arabidopsis roots [28, 32], we further impose that in the QC the WT BRAVO promoter must be ~ 0.2 times the value of the WOX5 promoter, i.e. *pB^WT^* (*x* = 0)/*pW^WT^* (*x* = 0) ~ 0.2. The parameter values in panels B and C (i.e. those corresponding to the red crossed in panels D and E) of Figure 2, Figure 3 and Figure 4 satisfy all these conditions, resulting in values ofthat qualitatively match the experimental observations in the *bravo* mutant.

### 4.6 Construction of realistic root layouts

In order to build two-dimensional realistic root layouts with which we can subsequently implement the corresponding reaction-diffusion equations (e.g. the Mixed model in next subsection), we start by taking a confocal image of a middle plane of the root with PI-stained cell walls and we apply a segmentation routine which divides the root at the pixel scale and into its constituent cellular regions and cell walls (Supplementary Figure 5). To do this, we make extensive use of the *scikit-image* collection of Python-based algorithms (*threshold_otsu, skeletonize* and *label*) [47]. In particular, we first define the cell boundaries with the *threshold_otsu* method, which transforms the original image into a thresholded binary image where only cell wall pixels remain. We then *skeletonize* to obtain a cell wall with a fixed width (2 pixels). We then apply the *label* function on the modified image to label distinct regions (each label constitutes the collection of pixels belonging to the particular region). We chose to define labels that enable to distinguish between cells and between the cell wall and the outside of the root as follows. All pixels within a cell have the same label, which is distinct from that of pixels in any other cell. With this routine, we can access each cell as an individual entity, and we can modify its properties as a whole (e.g. change the parameter values of the protein dynamics in all pixels of that cell). A single label is assigned to all the pixels in any cell wall. Thus all cell walls constitute a single, the same, entity. Another single region is defined by all pixels outside of the root.

The pixel grid is the spatial grid on which the dynamical equations of protein concentrations are settled. For the images used we have 1 pixel ≈ 0.5 *μ*m. We assign the same dynamical equations and parameter values in all pixels within each labelled region. These equations and parameter values can be distinct between regions. Specifically, the equations applied on the cell wall are distinct to those within cells, as described in the next subsection. The diffusion coefficient of a protein within all pixels of the cell wall is the same. For simplicity, the diffusion coefficient of a protein is set to be the same in all the cell regions (i.e. within any cell), but distinct from that in the cell wall. The only differences settled between different cells are on the protein production terms. Further details on the construction and implementation of the model in the realistic root layouts can be found in the SI Text.

### 4.7 Mixed model in a realistic root layout

In the Mixed model, WOX5 activates BRAVO through the protein *Z*, which is repressed by BRAVO. WOX5 negatively regulates its own production [28]. Both WOX5 and BRAVO bind to form an immobile and inactive complex. WOX5 and *Z* diffuse inside cells and between cells. For simplicity, no diffusion for BRAVO inside cells is settled. The equations for the rate of change of the concentrations of each type of protein across space *r* and time *t* are:

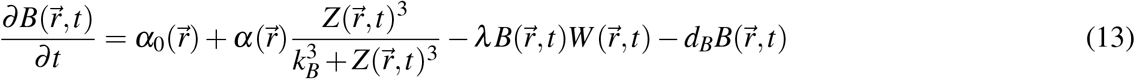

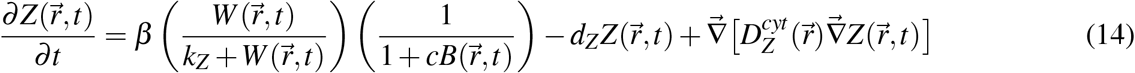

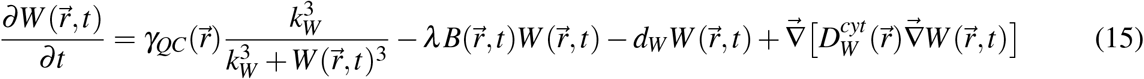

The GFP proteins produced under the promoters of BRAVO (*B_GFP_*), of WOX5 (*W_GFP_*) and of Z (*Z_GFP_*) have the same production rate as BRAVO, WOX5 and Z, respectively. All these GFP proteins have the same diffusion coefficient and all degrade linearly with the same degradation rate (i.e. that of GFP, *d_GFP_*). Hence, the equations for the dynamics inside cells of these GFP proteins are:

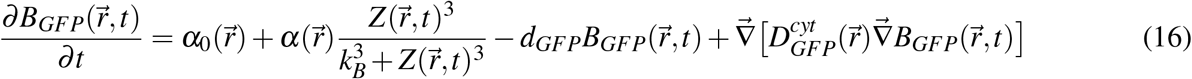

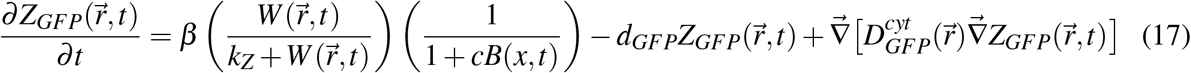

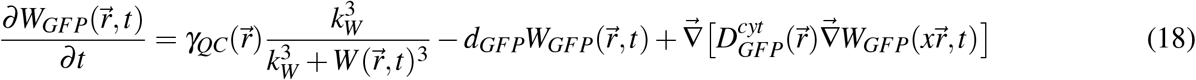

In these equations, 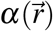 is the maximum strength of activation of BRAVO by *Z*, and is set to take the value *α* only in the cells shown in Supplementary Figure 6, being zero in all the other cells and regions. The threshold *k_B_* controls the levels of *Z* necessary to activate BRAVO. 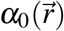 is the basal production rate of BRAVO, and takes the value *α*_0_ only in the pixels corresponding to the cells shown in Supplementary Figure 7 (being zero in all other regions). The factor *Z* is activated by WOX5 with maximum rate β, and activation threshold *k_Z_*. Parameter *c* sets the strength of the repression that BRAVO does on *Z*. 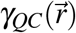 sets the maximal production of WOX5, and takes the value *γ_QC_* only in the regions corresponding to QC cells (Supplementary Figure 6), being zero in the remaining regions. *K_W_* sets the WOX5 concentration threshold to feed negatively back on its own production. The formation of the complex between BRAVO and WOX5, and the degradation of all factors are modelled as in Sections 4.1, 4.2, 4.3. The diffusion coefficient for each species within cells is indicated by superscript *cyt*. The diffusion terms take into account that across the whole root layout the diffusion coefficients are not homogeneous, since they take one value inside cells (superscript *cyt*) and another value in the cell walls (superscript *wall*).

We impose that inside cell walls only diffusion can happen whereas no reaction (production, degradation or binding), can occur, leading to the following equations:

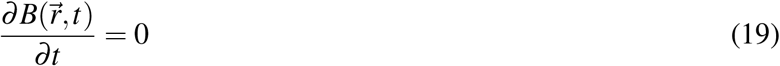

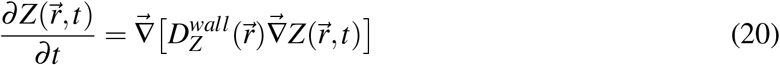

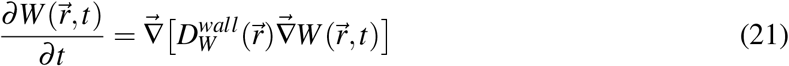

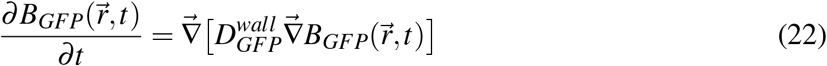

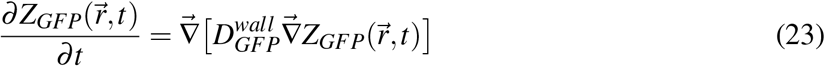

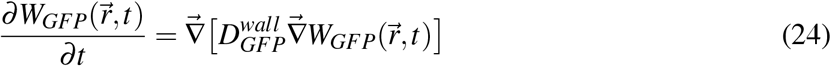

were the diffusion coefficient in the cell wall (denoted by superscript *wall*) is distinct to that inside cells. In Figure 5, *pBRAVO:GFP* corresponds to the variable 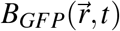 computed in the stationary state (i.e. the activity of the BRAVO promoter as seen through its GFP reporter), while 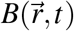 represents the BRAVO protein concentration, also computed in the stationary state. The analogous definitions for WOX5 and Z are used in Supplementary Figure 9.

All equations described up to this point correspond to the WT condition. To model the *bravo* mutant, the same equations, with the same parameter values are used, but the BRAVO protein concentration is set to zero (i.e. the mutant corresponds to Eqs. 14–18, 20–24 and 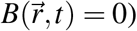. Notice that to model this mutant, the GFP reporter for BRAVO promoter, *B_GFP_*, is described with the same equations as in the WT (i.e. with Eqs.(16, 22)) and hence is not set to zero.

Figure 5 B and C show results of this same model (Eqs. 13, 15–24) except for the dynamics of *Z* within cells which, instead of Eq. (14), is set as:

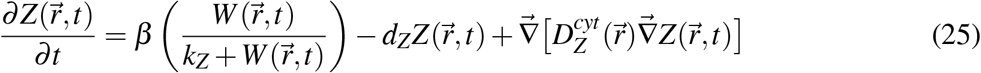

Notice that this is the same Eq.(14) except for the repression term by BRAVO, which here is not present. In addition, in Figure 5 B, *Z* does not diffuse and hence 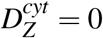 and 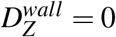. In Figure 5D the equations of the Mixed model are used with *λ* = 0.

### 4.8 Numerical implementation of the mathematical models

To find the stationary distributions of the 1D models described in Sections 4.1, 4.2 and 4.3, we first reduce the system of differential equations to second order ordinary differential equations for the diffusible variables. To do this, we first set to zero the equation for the non-diffusible variables and substitute the result on the other equations. The remaining system can be cast as a boundary value problem which we solve numerically by using the *solve_bvp* routine embedded in the Python-based *SciPy* library [48]. In all these calculations, we discretize the 1D Laplacian term as 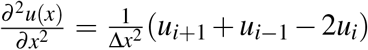, being *i* an index such that *x* = *i*Δ*x*, and use a spatial step size of Δ*x* = 0.05 a.u., in a domain of *x* ∈ [–600,600] a.u. We use a QC size of *L_QC_* = 15 a.u., and an equally long VI region. With this methodology, the explicit time-dependence of the variables is not computed, but only their stationary state. To double-check the validity of our solutions, we also simulated the whole temporal dynamics of the equations with a forward Euler method, obtaining the same stationary solutions as with the *solve_bvp* method, thus confirming the results (data not shown).

To simulate the dynamics of the Mixed model (and its modifications) in the realistic root layout, we solve the corresponding reaction-diffusion equations with heterogeneous diffusion coefficients with a forward-time central-space scheme (FTCS) where time is discretized in steps of size Δ*t* = 0.1 a.u., (so that after *k* steps, *t_k_* = *k*Δ*t*) and space is discretized in steps of size (Δ*x* =1, Δ*y* = 1) pixels, so that (*x_i_,y_i_*) = (*i*Δ*x*, *j*Δ*y*), with a lattice size of (*L_x_,L_y_*) = (228,448) pixels for WT root and (*L_x_,L_y_*) = (231,448) pixels for *bravo* mutant root. For the images used, 1 pixel≈0.5μm. We run the simulations up to *t* = 4000 a.u., and take this time point as corresponding to the stationary state. All the diffusion coefficients outside the root layout are set to be zero, restricting the domain of the equations to the root’s interior. Additionally, this condition automatically implements reflecting boundary conditions at the root borders.

## 5 Author contributions

JM and MI designed the research with the help of IBP, NB and AICD. JM and MI formulated the mathematical models. JM performed the numerical simulations. All authors analyzed the data. JM and MI wrote the manuscript with the help of IBP, NB and AICD.

## 6 Acknowledgements

M.I. and J.M. acknowledge support from grant PGC2018-101896-B-I00 funded by MCIN/AEI/10.13039/ 501100011033/ FEDER “Una manera de hacer Europa”, and from the Generalitat de Catalunya through Grup de Recerca Consolidat 2017 SGR 1061. A.I.C-D. is a recipient of a BIO2016-78955 grant from the Spanish Ministry of Economy and Competitiveness and a European Research Council, ERC Consolidator Grant (ERC-2015-CoG–683163). J.M. acknowledges BES-2016-078218 funded by MCIN/AEI /10.13039/501100011033 and FSE “El FSE invierte en tu futuro”. I.B-P. is funded by the FPU15/02822 grant from the Spanish Ministry of Education, Culture and Sport; N.B. by the FI-DGR 2016FI-B 00472 grant from the AGAUR, Generalitat de Catalunya; CRAG is funded by “Severo Ochoa Programme” from Centers of Excellence in R&D 2016-2019 (SEV-2015-485 0533).

## Supplementary Information Text

## 1 Derivation of the models

The equations used in all models come from an approximation where complex formation and mRNA dynamics is very fast compared to the dynamics of proteins. We exemplify this approximation with the immobilization by sequestration mechanism, but the same procedure has been applied to obtain the set of equations of all the other models.

By explicitly considering the mRNA of BRAVO (m_B_) and WOX5 (*m_W_*), and the complex formed by the binding of BRAVO and WOX5 proteins (*C*), the model equations for all these variables and for BRAVO (*B*) and WOX5 (*W*) proteins in the immobilization by sequestration mechanism are:

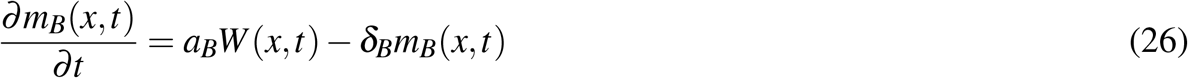

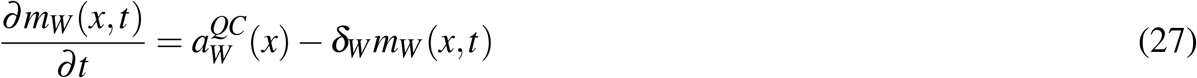

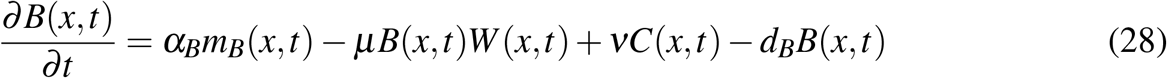

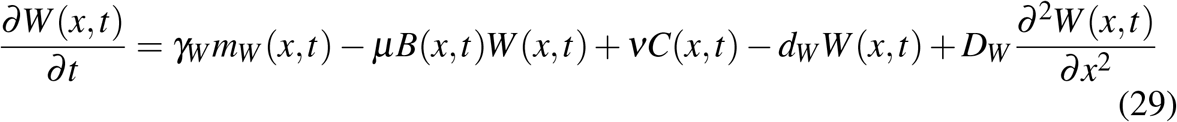

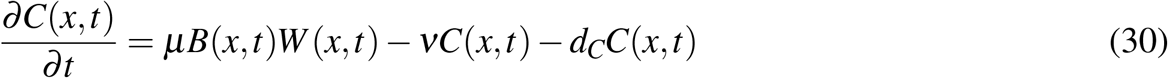

where 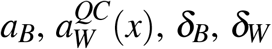 are the mRNA synthesis and degradation rates of BRAVO and WOX5 mRNAs, respectively. The superscript and explicit spatial dependence in 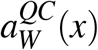 indicates that the mRNA of WOX5 is only produced in the QC region. *α_B_* and *γ_W_* represent the rates of mRNA translation into BRAVO and WOX5 proteins, respectively, and *μ* and *ν* are the rates of protein-protein binding and unbinding. Finally, the complex can be degraded with rate *d_C_*. The rest of the parameters have already been defined in Methods. If the dynamics of the mRNAs and of complex formation are very fast (by setting their corresponding time derivatives to zero), we obtain:

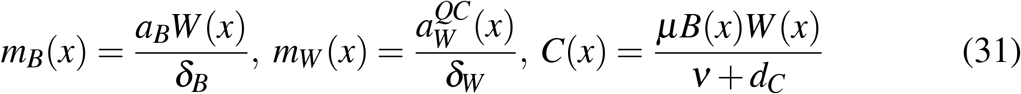

Substituting these relations to the equations for *B* and *W*, it results into:

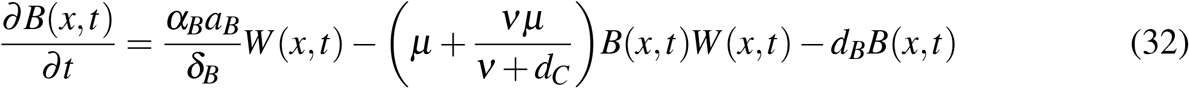

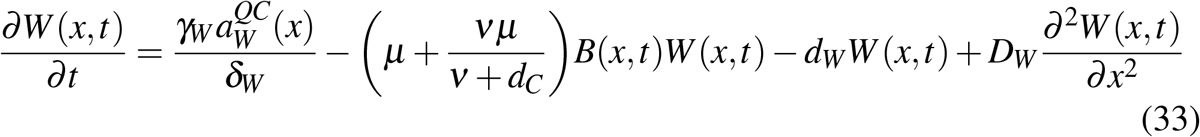

These are the equations of the immobilization by sequestration model used in the main text (Eq. 1 and 2) when the following definitions of parameters are applied: 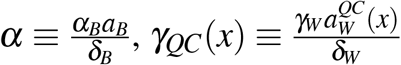 and 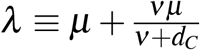.

Notice that since we only analyse the stationary state of the system, these quasi-steady state approximations do not affect the final result of the spatial profiles.

### 1.1 Stationary profile of WOX5 in the immobilization by sequestration model

In the stationary state, the previous equations (32) and (33) of the immobilization by sequestration model can be reduced to the following second order ODE:

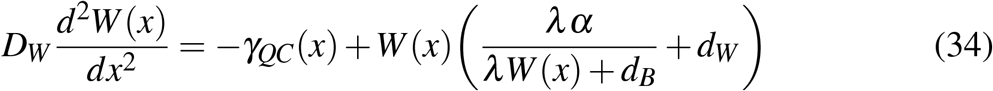

and the BRAVO profile is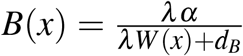 can be interpreted as a defusing molecule *W* produced at a source *γ_QC_*(*x*), with a *nonlinear* higher degradation rate, given by

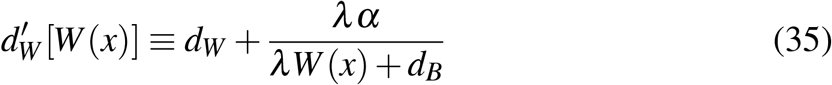

where the second term comes from the binding between BRAVO and WOX5. This non-linear degradation implies that in the immobilization by sequestration model and outside the source region, the stationary spatial profile of *W*(*x*) in the WT is not exponential, but decays spatially more abruptly. In contrast, when modeling the *bravo* mutant, the same ODE applies but with a linear degradation, 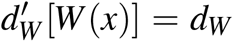 and hence the profile of *W*(*x*) outside the source region is exponential in this mutant.

## 2 Immobilization by sequestration with an additional sequestra-tor

In Supplementary Figure 2 we show the effect of an additional protein (hereafter named sequestrator, *S*) which can bind BRAVO and WOX5 separately. The equations corresponding to this model are:

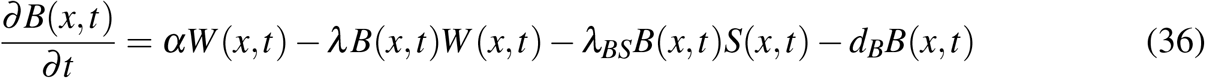

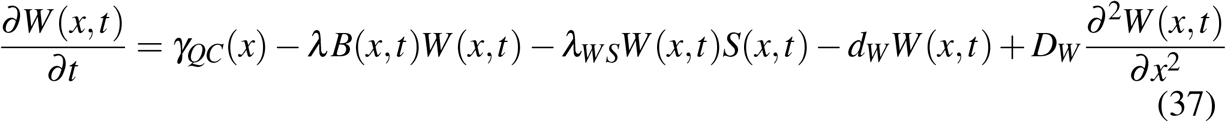

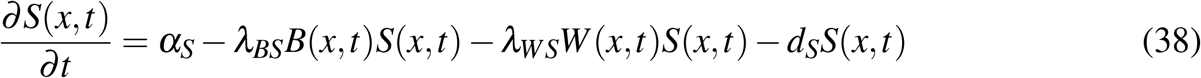

where *S*(*x, t*) is the concentration of the protein *S* across space and time, the new parameters *α_S_* and *d_S_* represent the production and degradation of *S* proteins, and *λ_BS_*, *λ_WS_* denote the complex formation rates between *S* and BRAVO and *S* and WOX5, respectively.

## 3 Simulations in a realistic root layout

### 3.1 Space discretization for diffusion with heterogeneous coefficients

In the realistic root layout, we simulate a reaction-diffusion equation with non-homogeneous diffusion coefficients, that is, diffusion is explicitly dependent on space. In our case, the value of these coefficients depend on whether the spatial position corresponds to the interior of a cell or to the cell wall, with respective diffusion coefficients of *D^cyt^* and *D^wall^*. These can be encompassed into a single, spatially-dependent coefficient *D*(*x, y*), where *x, y* carry the information of the positions within the root layout:

*D(x, y) = D^cyt^* for *x, y*∈cells and *D(x, y) = D^wall^* for *x, y*∈cell wall.

Then, for a given variable *u*(*x,y*) (i.e. the concentration of one of the proteins), we discretize the spatial term 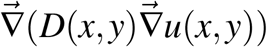 as done in [49], namely:

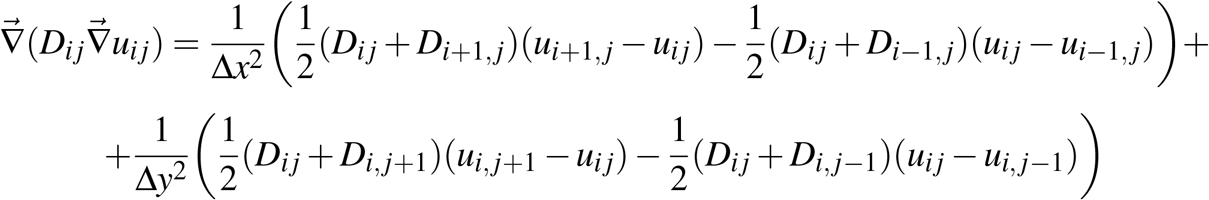

where *D_ij_* = *D*(*x* = *i*Δ*x,y* = *j*Δ*y*) and *u_ij_* = *u*(*x* = *i*Δ*x,y* = *j*Δ*y*). Thus the values of indexes *i*, *j* specify the value of the diffusion coefficient, whether it is *D^cyt^* or *D^wal1^*.

### 3.2 Reaction-diffusion model with only WOX5

Supplementary Figure 8 shows the stationary results when WOX5 is produced only at QC cells, degrades and diffuses, in the absence of any other regulation. In this case only the dynamics for WOX5 and the GFP reporter its promoter are simulated, according to the following equations inside the cells:

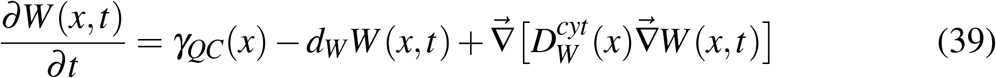

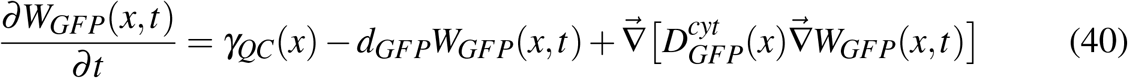

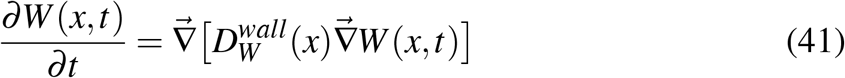

and in the cell wall:

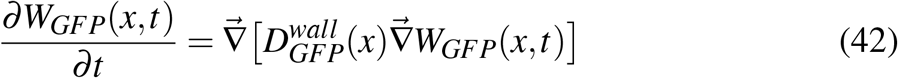

Notice that for these variables, these are the same equations as those of the Mixed model but with *λ* = 0.

## Supplementary Figures

**Figure 6: Supp. Fig. 1.**
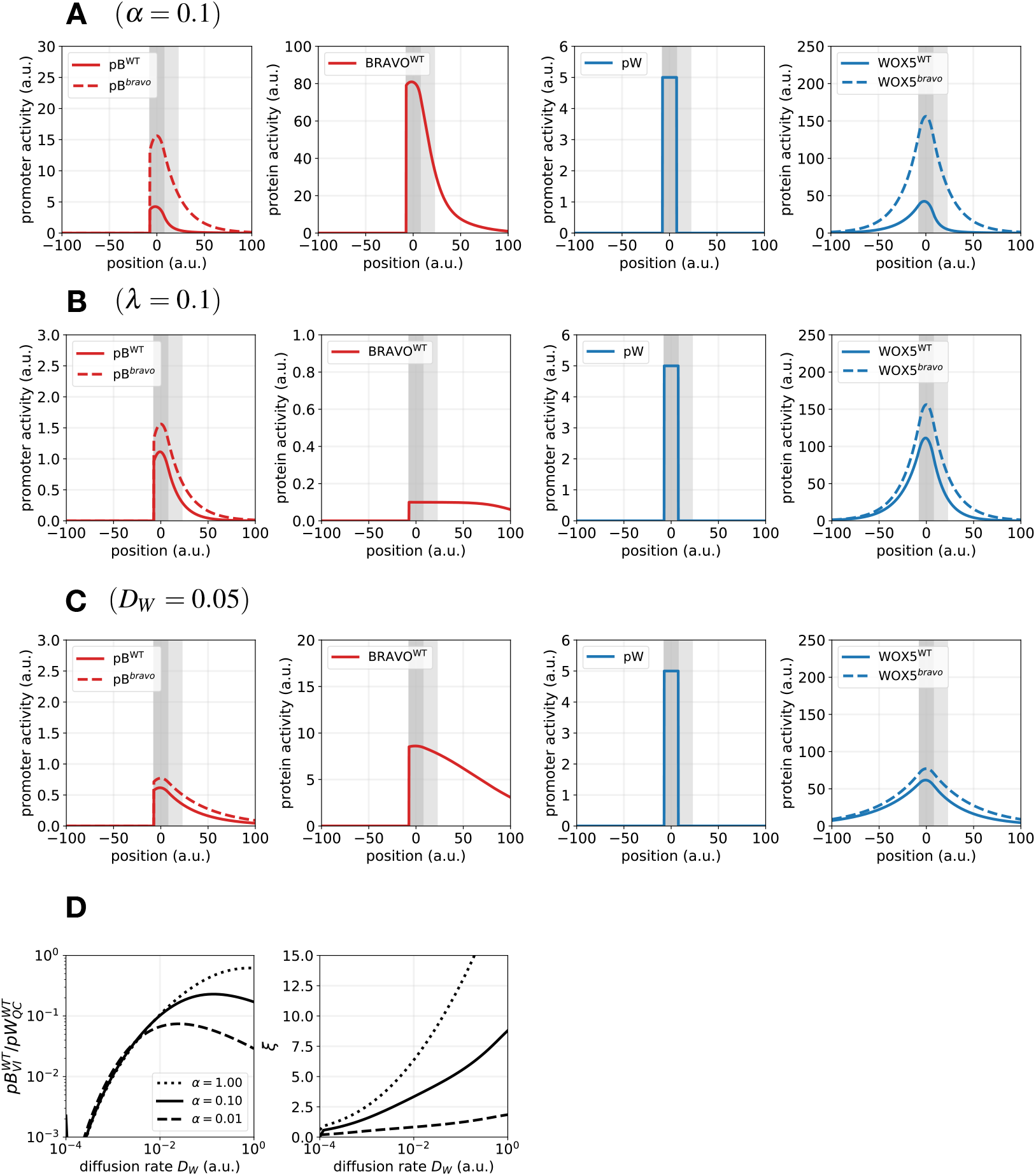
Results of the Immobilization by sequestration model for different parameter values. Stationary profiles of *pB*(*x*), *B*(*x*), *pW*(*x*) and *W*(*x*) in WT (continuous lines) and in the *bravo* mutant (dashed lines), for the same parameter values as in Figure 2 except for one: **A)** *α* = 0.1, **B)** *λ* =0.1 and **C)** *D_W_* = 0.05. Accordingly, results in A, B and C, when compared to Figure 2, depict the effect of (A) higher BRAVO production, (B) stronger binding or (C) higher WOX5 diffusion. **A)** For this higher production rate, *pB* in the WT is mostly at the QC (with similar levels to *pW*) and nearly absent in the VI. This strong confinement is not compatible with real expressions in Arabidopsis roots. **B)** This higher complex formation rate has a strong impact on the levels of free BRAVO, which are very small due to higher sequestration by WOX5. **C)** For this higher WOX5 diffusion coefficient, WOX5 is at high concentration across a very broad region above the VI. Consequently *pB*(*x*) is also very spanned, which is not realistic when compared to expressions in Arabidopsis roots. *pB* value at the QC is lower than in Figure 2. **D)** The effect of WOX5 diffusion coefficient on the quantities 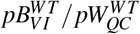 and *ξ*, for different values of *α*. A non-zero value of WOX5 diffusion is needed to induce an expansion of *pB*(*x*) in the *bravo* mutant. Yet, too large diffusion coefficients reduce the level of *pB* expression at the VI and QC.

**Figure 7: Supp. Fig. 2.**
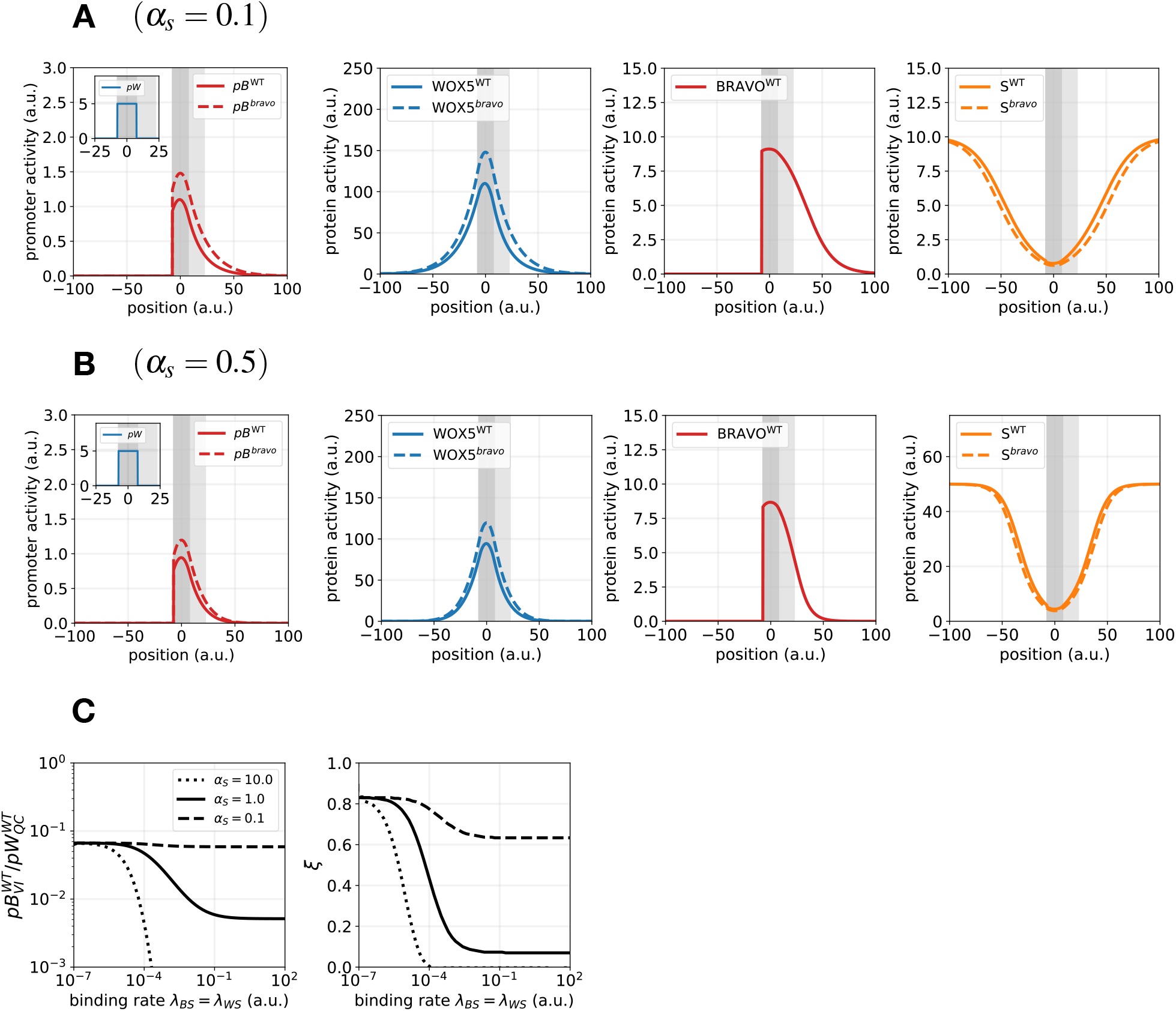
Immobilization by sequestration model with an additional sequestrator *S*. All common parameter values as in Figure 2. **A,B)** Stationary profiles of *pB*(*x*), *B*(*x*), *pW*(*x*), *W*(*x*) and *S*(*x*) for two different values of the production of the additional sequestrator (A) *α_S_* = 0.1 and (B) *α_S_* = 0.5. The rates of binding between S and BRAVO and between S and WOX5 are the same as that between BRAVO and WOX5, *λ_BS_*= *λ_WS_* = *λ_W_B* =. **C)** 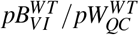 and *ξ* as a function of the binding rates *λ_BS_* = *λ_WS_*, for three different *α_S_* values. An increase in *α_S_*, *λ_BS_* and *λ_WS_* values weakens *pB*(*x*) expansion. The other parameter values of *S* dynamics are detailed in Table 2.

**Figure 8: Supp. Fig. 3.**
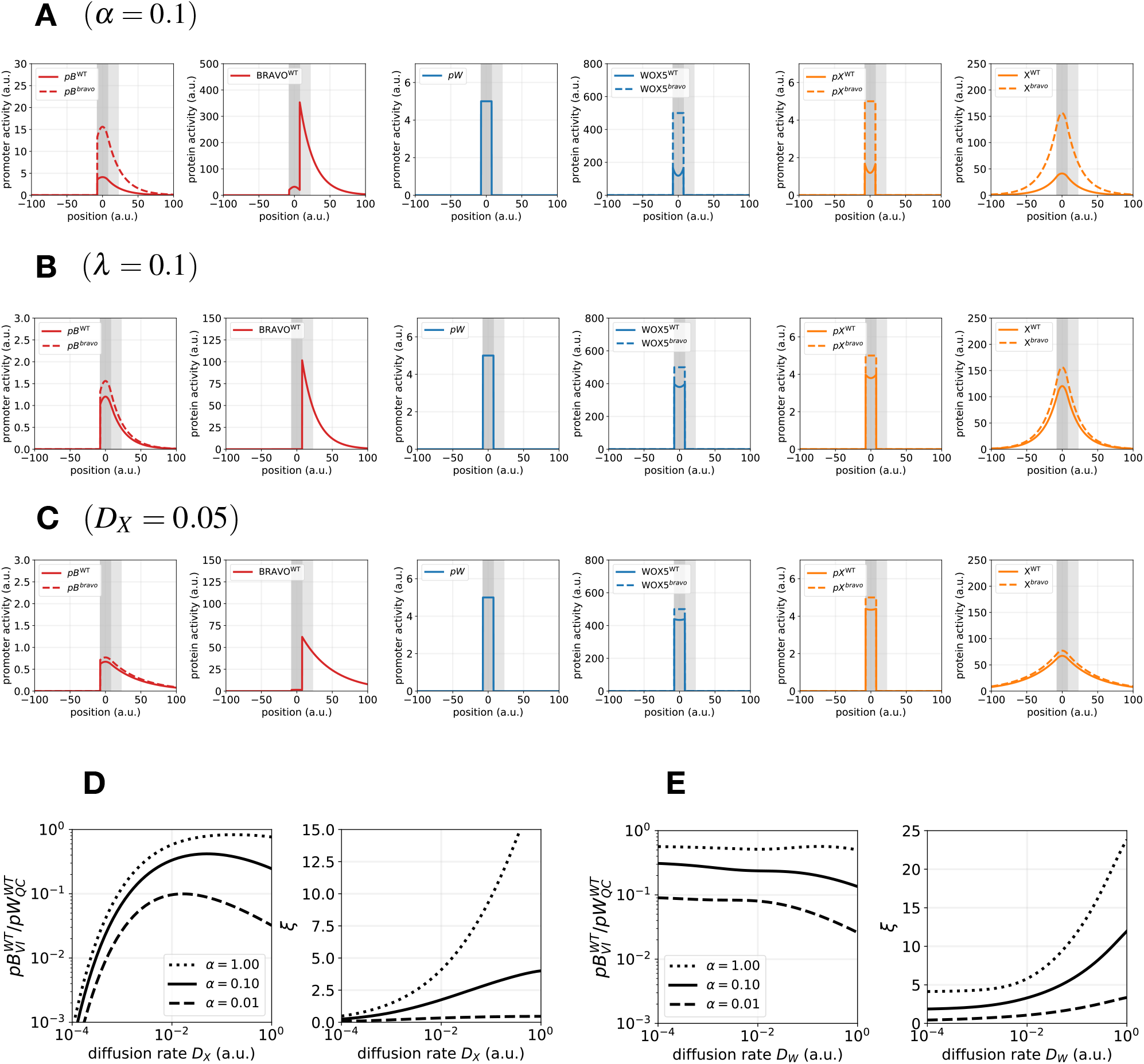
Results of the Attenuation by sequestration model for different parameter values. Stationary profiles of *pB*(*x*), *B*(*x*), *pW*(*x*), *W*(*x*), *pX*(*x*) and *X*(*x*) in WT (continuous lines) and in the *bravo* mutant (dashed lines), for the same parameter values as in Figure 3 except for one: **A)** *α* = 0.1, **B)** *λ* = 0.1 and **C)** *D_X_* = 0.05. Accordingly, results in A, B and C, when compared to Figure 3, depict the effect of (A) higher BRAVO production, (B) stronger binding or (C) higher *X* diffusion. **A)** For this higher production rate, *pB* in the WT at the QC has similar levels to *pW*, a situation which is not compatible with real expressions in Arabidopsis roots. The bump in the profile of *B*(*x*) is a direct consequence of sequestration only happening in the QC, as in this model WOX5 does not diffuse. **B)** For this higher complex formation rate, BRAVO is nearly absent from the QC, being all sequestered by WOX5. The profiles of *pB*(*x*) are very similar to the ones in Figure 3. **C)** For this higher *X* diffusion coefficient, *pB*(*x*) is very spanned above the VI, which is not realistic when compared to expressions in Arabidopsis roots. *pB* value at the QC is lower than in Figure 3. **D)** Effect of *X* diffusion coefficient on the quantities 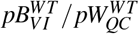 and *ξ*, for different values of *α*. In panels A-D there is no diffusion of WOX5. **E)** Effect of *WOX*5 diffusion coefficient on the quantities 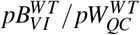 and *ξ*, for different values of *α*. The diffusion of WOX5 promotes expansion of *pB*(*x*) in the *bravo* mutant but drives fainter *pB* values at the VI (and QC) in the WT.

**Figure 9: Supp. Fig. 4.**
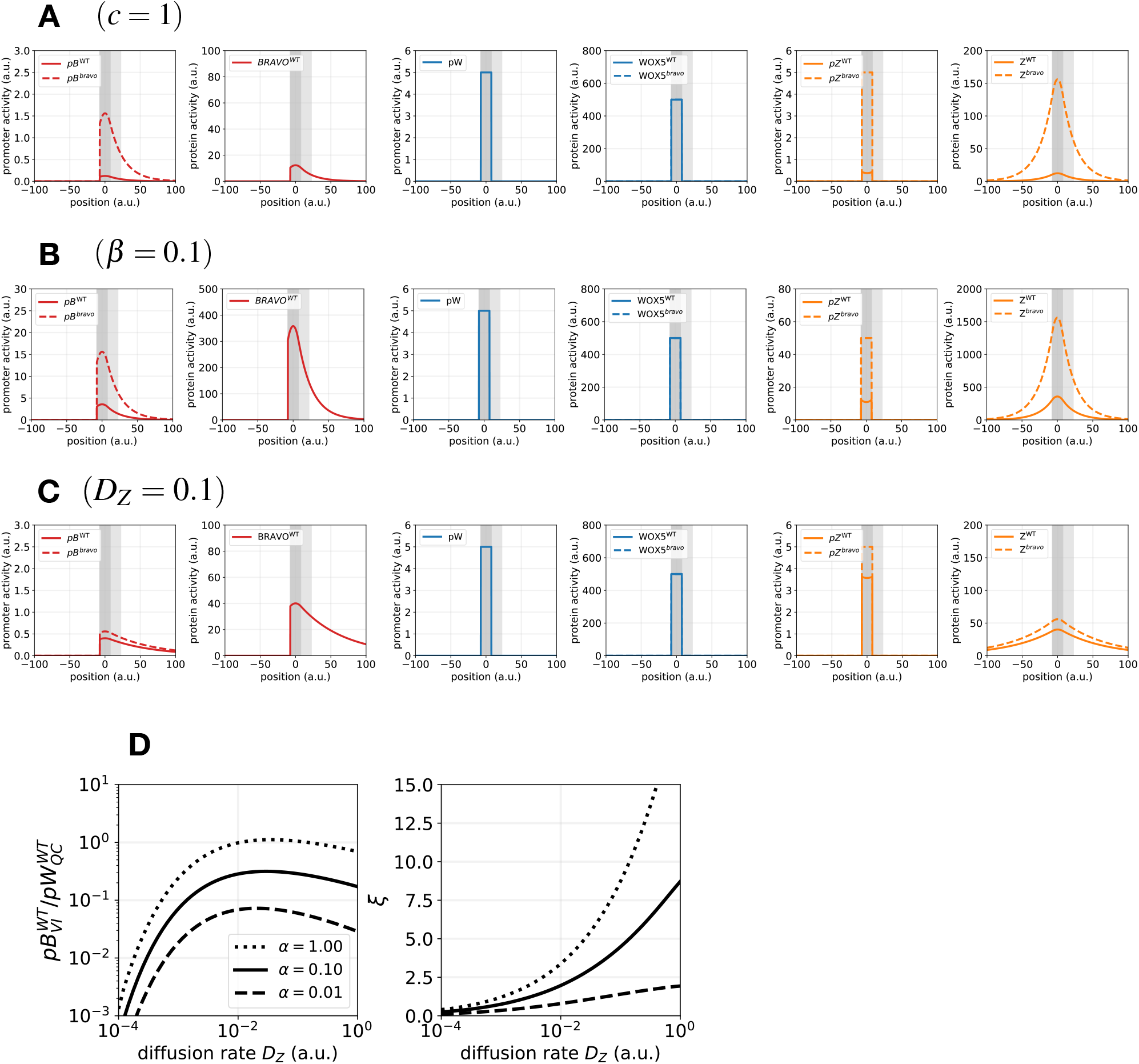
Results of the Repression model for different parameter values. Stationary profiles of *pB*(*x*), *B*(*x*), *pW*(*x*), *W*(*x*), *pZ*(*x*) and *Z*(*x*) in WT (continuous lines) and in the *bravo* mutant (dashed lines), for the same parameter values as in Figure 4 except for one: **A)** *c* = 1, **B)** *β* = 0.1 and **C)** *D_Z_* = 0.1. Accordingly, results in A, B and C, when compared to Figure 4, depict the effect of (A) higher repression strength, (B) higher production rate of *Z* or (C) higher *Z* diffusion. **A)** For this higher repression strength, *pB* in the WT is very low, and increases and spans very dramatically in the *bravo* mutant, which is not compatible with experimental data (Figure 1, [28]). **B)** For this higher production rate of *Z*, *pB* in the WT is very high, and increases very dramatically in the *bravo* mutant, which is not compatible with experimental data. **C)** For this higher *Z* diffusion coefficient, *pB*(*x*) is very spanned above the VI, which is not realistic when compared to expressions in Arabidopsis roots. *pB* value at the QC is lower than in Figure 3. **D)** Effect of *Z* diffusion coefficient on the quantities 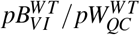 and *ξ*, for different values of *α*. Strong diffusion of *Z* promotes expansion of *pB*(*x*) in the *bravo* mutant but drives fainter *pB* values at the VI (and QC) in the WT.

**Figure 10: Supp. Fig. 5.**
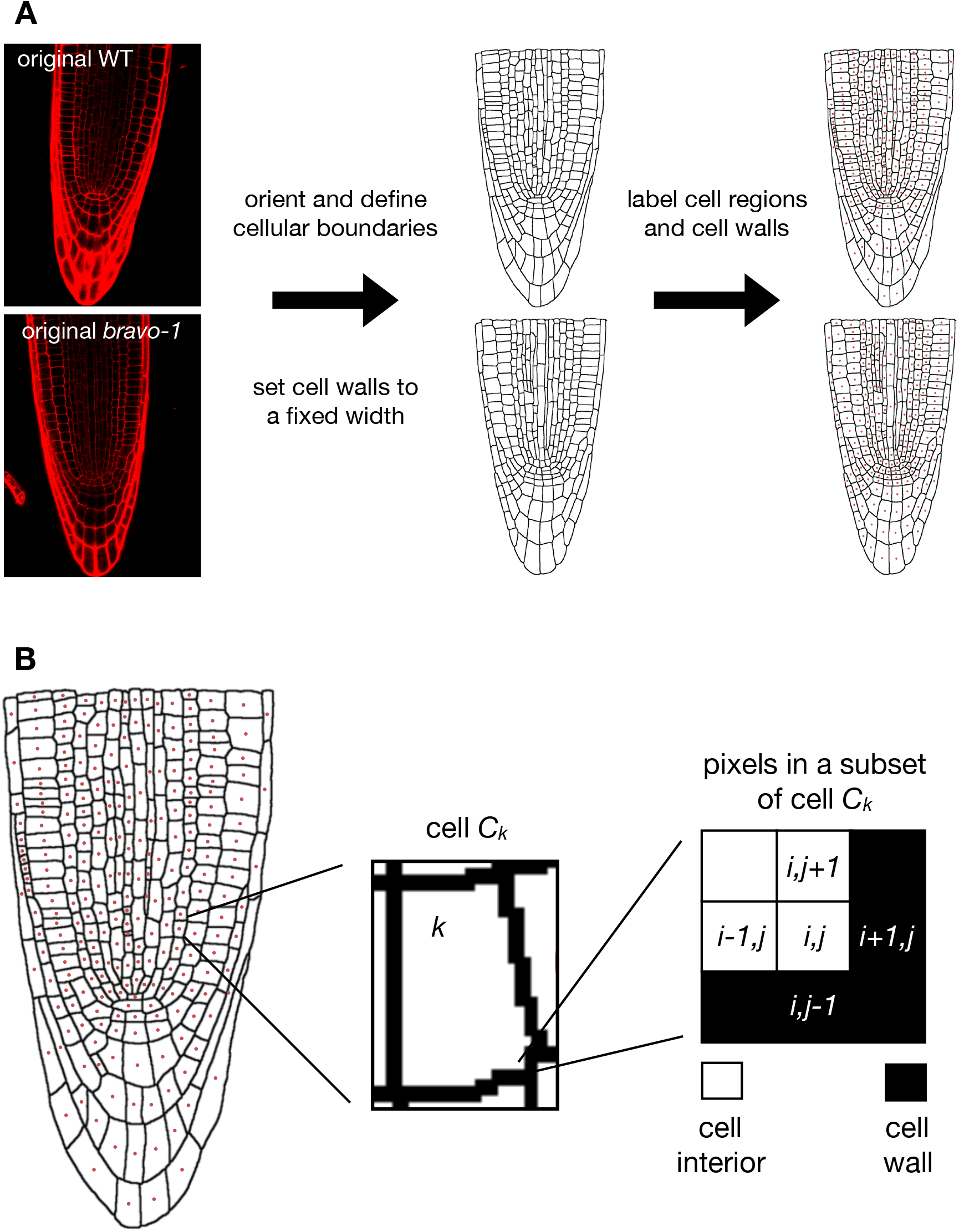
Construction of a realistic root layout. **A)** We select two root tips, representative for WT and *bravo-2* mutants, from a confocal image of the root where cell walls are PI-stained (red). We first orient and re-scale the roots so that both can be compared. As a result proportions are slightly modified from the original image. This initial step is optional. We then reset a fixed width for the cell walls (2 pixels in the simulation). Subsequently, we label the images to define each cellular region (marked as red dots located at the centroid of each cell). **B)** Cell *C_k_* is the cell that contains the pixels with label *k* (which define the cell’s interior, white). It is surrounded by pixels corresponding to cell walls (black). The spatial position of a pixel is denoted by two indexes, *i* and *j*. On this discretized grid we implement the corresponding reaction-diffusion equations.

**Figure 11: Supp. Fig. 6.**
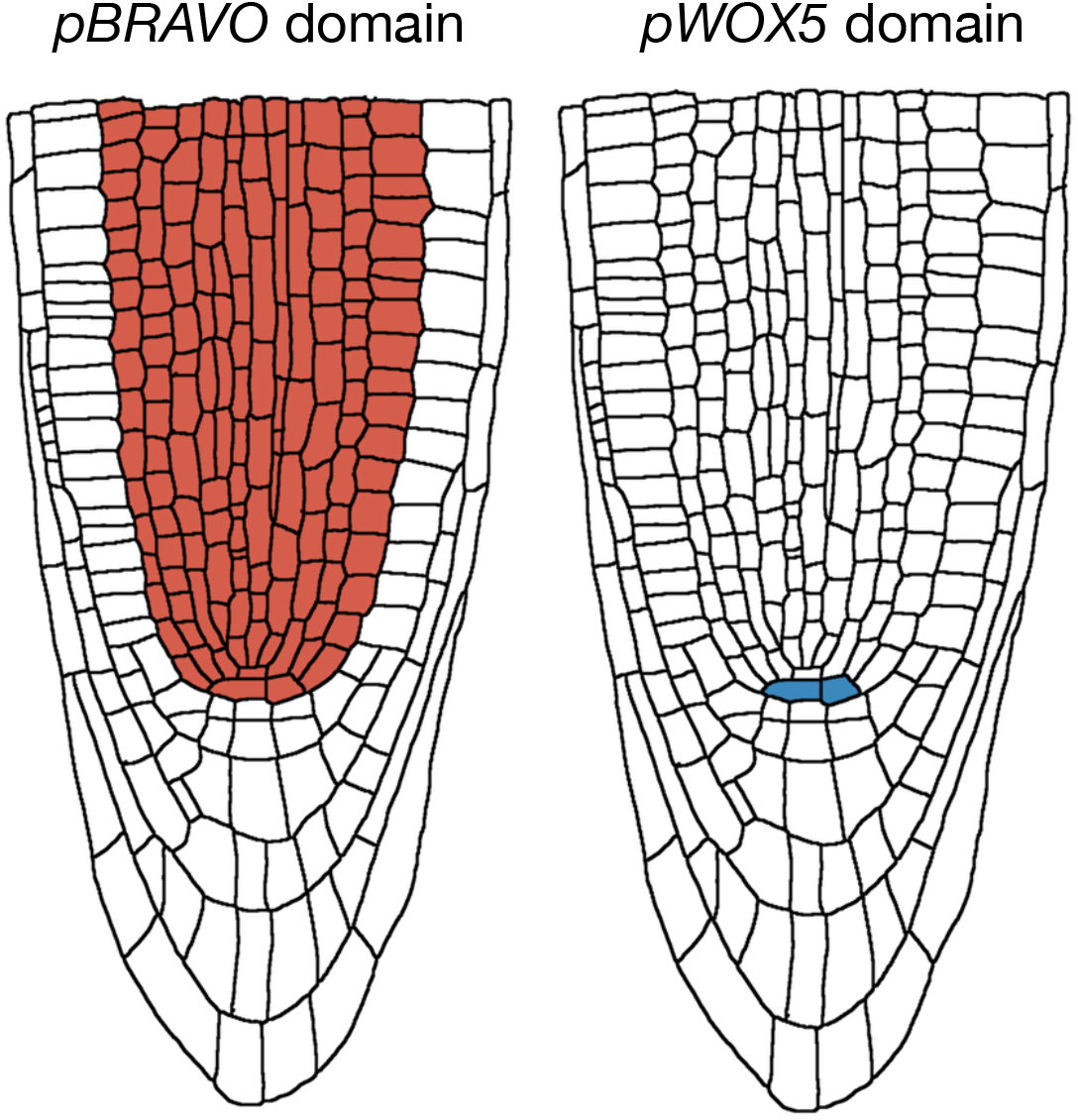
Cellular domains where production of BRAVO and WOX5 is enabled in the simulations. The production terms of BRAVO and WOX5 are restricted to the red and blue domains respectively (these regions are denoted as the pBRAVO and pWOX5 domains, since they indicate where the promoter of BRAVO and WOX5 can have an activity). The GFP reporters of their promoters are produced only in these same domains. In this way, we only allow *B* and *BGFP* to be activated by *Z* in the QC, vasculature, cortex and endodermis, while *W* and *W_GFP_* are only produced at the QC. This does not prevent WOX5 and GFP proteins to be present in other tissues, which they can reach through diffusion.

**Figure 12: Supp. Fig. 7.**
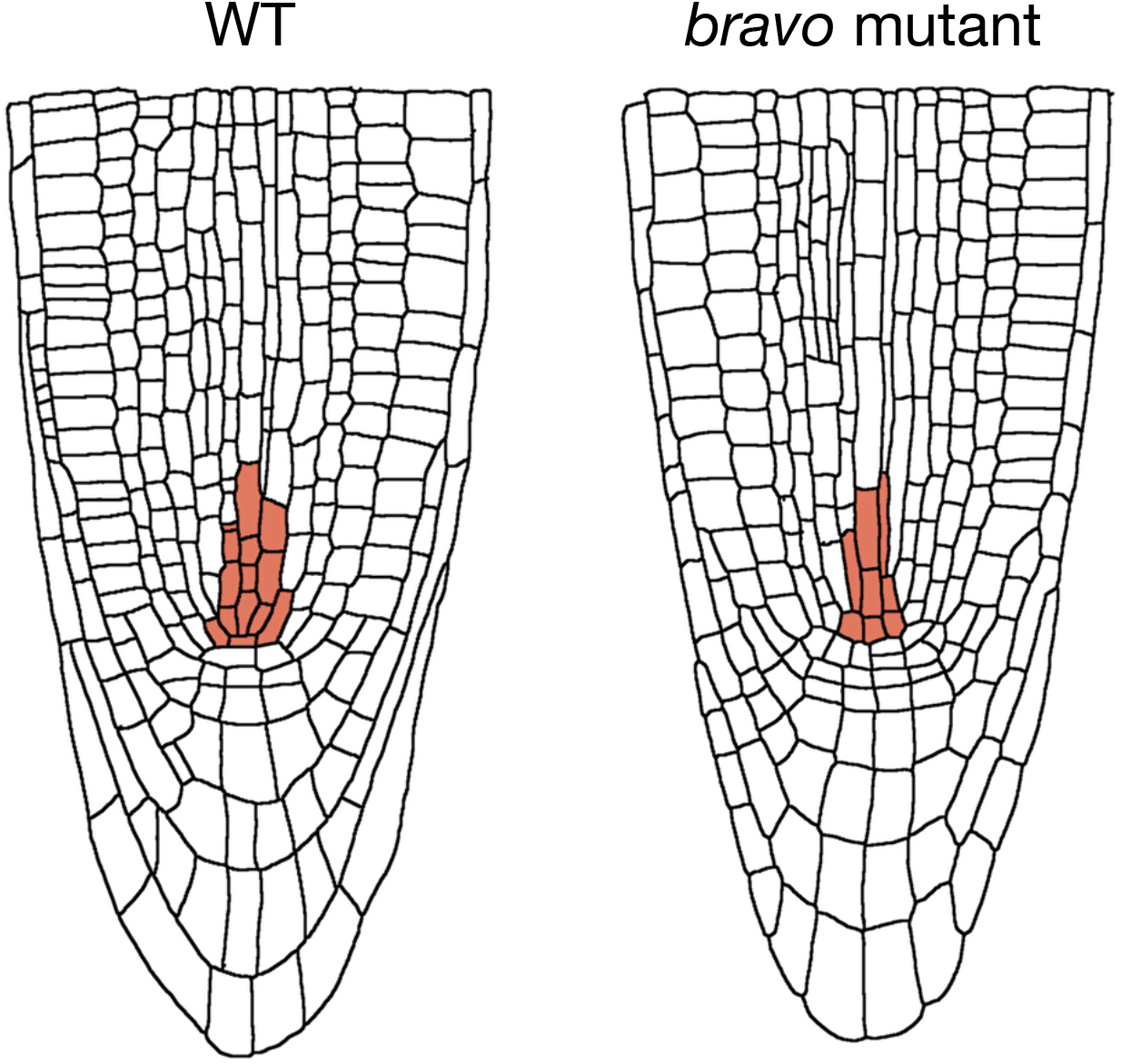
Domains of basal production of BRAVO. In the simulations using realistic root layouts, BRAVO is assumed to have basal levels of production only in those first cells of the vasculature colored in red. This consideration comes from the fact that in double *bravo wox5* mutants, basal expression of *pBRAVO:GFP* can still be observed [28], with a spatial pattern similar to the one shown in red in the figure. The GFP reporter of BRAVO promoter is set also to have a basal activity in the red domains only. Since WT and *bravo* mutant roots differ in morphology, the specific cells that are set to have basal BRAVO production are slightly different in each root, as depicted.

**Figure 13: Supp. Fig. 8.**
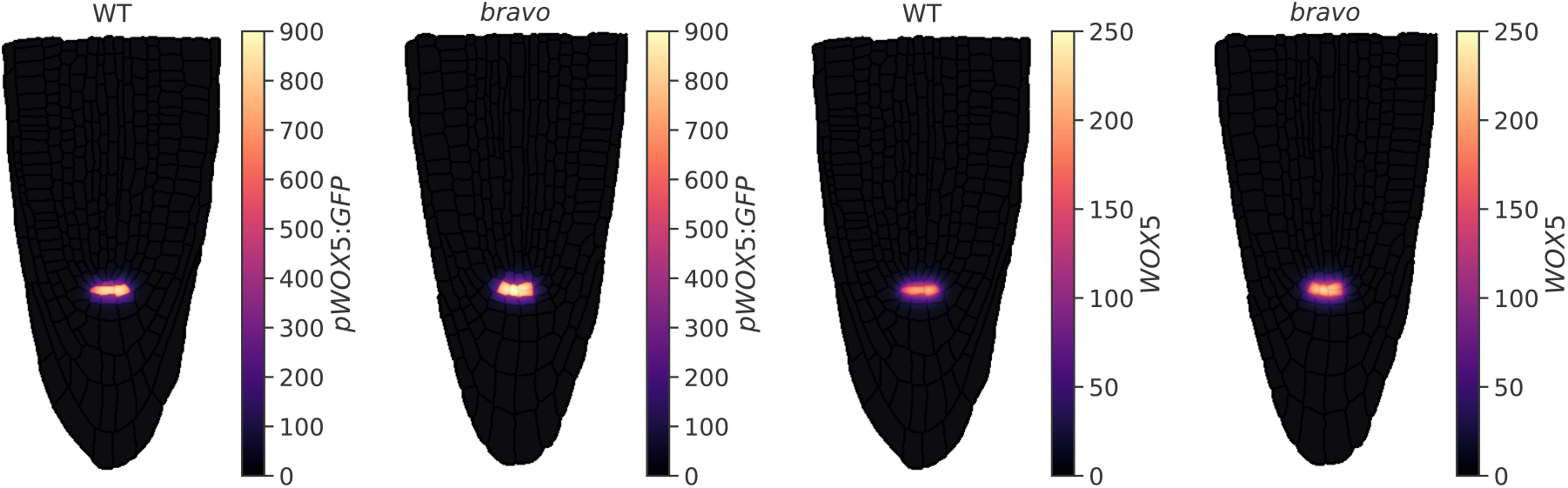
Concentration of *pWOX5:GFP* (*W_GFP_*) and *WOX5* (*W*) in the absence of other regulatory factors. Stationary patterns in the WT and in the *bravo* mutant when WOX5 is produced only at the QC, degrades and diffuses as in the Mixed Model, and no other molecule (e.g. BRAVO) is present. The model used is detailed in SI Text. All parameter values of WOX5 dynamics as in Figure 5. This WOX5 diffusion coefficient allows WOX5 to reach only with visible concentration the cells adjacent to the QC. Concentrations shown in this figure are obtained by running the simulations up to a time *t* = 3000 a.u., at which the stationary state is already reached.

**Figure 14: Supp. Fig. 9.**
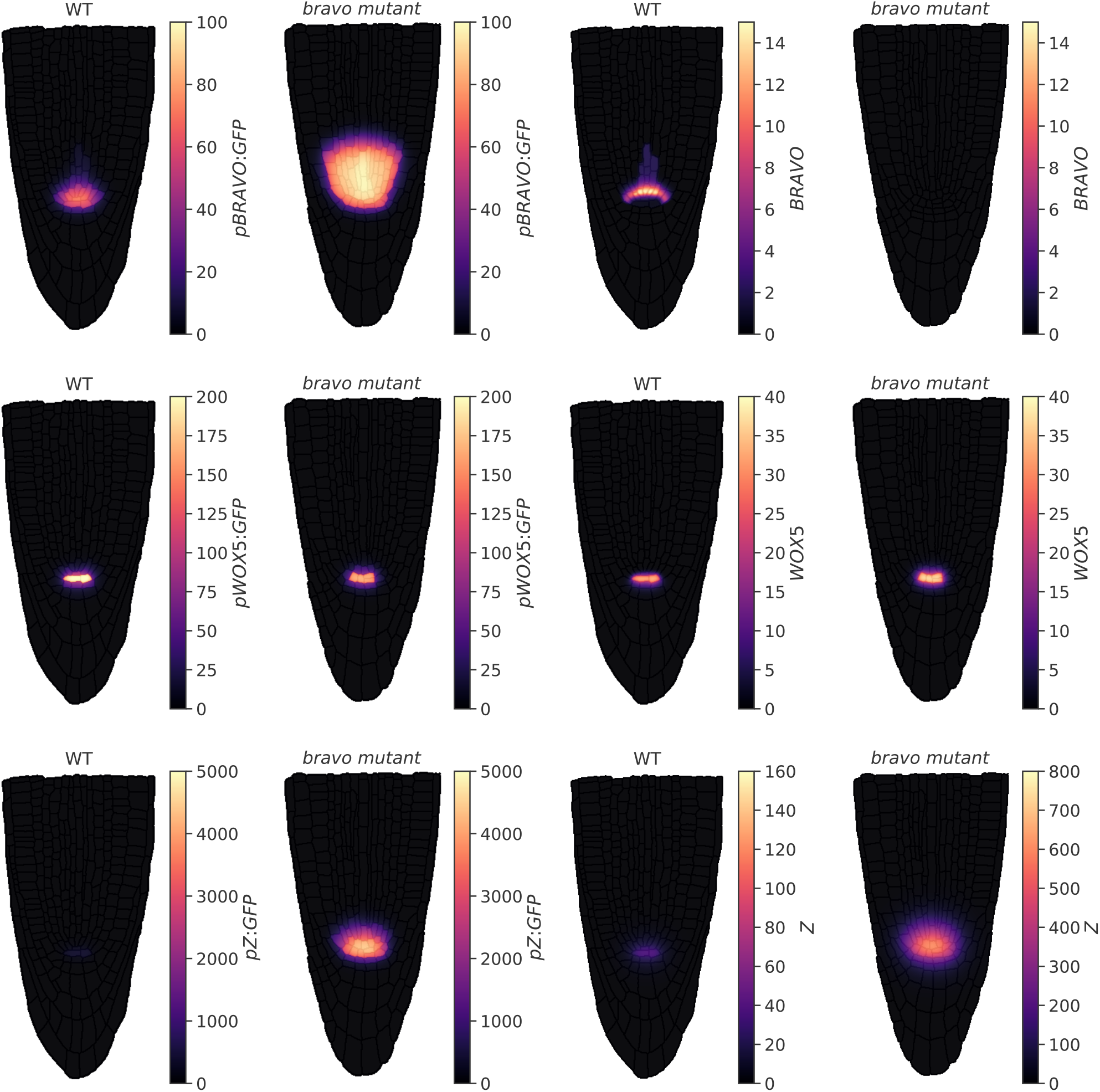
Stationary profiles of all proteins and GFP reporters of promoters corresponding to the case of Figure 5A (Mixed Model). The results are shown for the WT and the *bravo* mutant. The first two panels correspond to the same results shown in Figure 5A.

## Supplementary Tables

**Table 1:** Default parameters of the immobilization by sequestration (IS), attenuation by sequestration (AS) and repression (R) models. All parameter values are indicated in arbitrary units.

| Parameter | Description | IS | AS | R |
| --- | --- | --- | --- | --- |
| $\alpha$ | Protein production rate of BRAVO | 0.01 | 0.01 | 0.01 |
| $\gamma_{QC}$ | Protein production rate of WOX5 | 5 | 5 | 5 |
| $\beta$ | Protein production rate of $X$ | – | 0.01 | 0.01 |
| $c$ | Threshold of $Z$ repression by BRAVO | – | – | 0.01 |
| $\lambda$ | Binding rate between BRAVO and WOX5 | 0.001 | 0.001 | 0.001 |
| $d_B$ | Degradation rate of BRAVO proteins | 0.01 | 0.01 | 0.01 |
| $d_W$ | Degradation rate of WOX5 proteins | 0.01 | 0.01 | 0.01 |
| $d_X$ | Degradation rate of $X$ proteins | – | 0.01 | – |
| $d_Z$ | Degradation rate of $Z$ proteins | – | – | 0.01 |
| $D_W$ | Diffusion coefficient of WOX5 proteins | 0.01 | – | – |
| $D_X$ | Diffusion coefficient of $X$ proteins | – | 0.01 | – |
| $D_Z$ | Diffusion coefficient of $Z$ proteins | – | – | 0.01 |

**Table 2:** Additional parameters for the immobilization by sequestration model with an additional sequestrator (ISAS). The remaining parameters are the same as in the original immobilization by sequestration model. All parameter values are indicated in arbitrary units.

| Parameter | Description | ISAS |
| --- | --- | --- |
| $\alpha_S$ | Protein production rate of $S$ | 0.1 |
| $d_S$ | Degradation rate of $S$ proteins | 0.01 |
| $\lambda_{BS}$ | Complex formation rate between BRAVO and $S$ | 0.001 |
| $\lambda_{WS}$ | Complex formation rate between WOX5 and $S$ | 0.001 |

**Table 3:** Default parameters of the mixed model in the realistic root layout. All parameter values are indicated in arbitrary units of concentration, time and space.

| Parameter | Description | Value (a.u.) |
| --- | --- | --- |
| $\alpha_0$ | Basal production rate of BRAVO proteins | 0.01 |
| $\alpha$ | Regulated production rate of BRAVO proteins | 0.1 |
| $\gamma_{QC}$ | Production rate of WOX5 proteins | 2 |
| $\beta$ | Production rate of Z proteins | 6 |
| $k_B$ | Threshold of BRAVO activation by Z | 10 |
| $k_Z$ | Threshold of Z activation by WOX5 | 1 |
| $k_W$ | Threshold of WOX5 self-repression | 20 |
| $c$ | Threshold of Z repression by BRAVO | 10 |
| $\lambda$ | Binding rate between BRAVO and WOX5 | 0.005 |
| $d_B$ | Degradation rate of BRAVO proteins | 0.005 |
| $d_W$ | Degradation rate of WOX5 proteins | 0.005 |
| $d_Z$ | Degradation rate of Z proteins | 0.005 |
| $D_W^{cyt}$ | Diffusion coefficient of WOX5 proteins in the cytoplasm | 0.2 |
| $D_W^{wall}$ | Diffusion coefficient of WOX5 proteins in cell walls | 0.1 |
| $D_Z^{cyt}$ | Diffusion coefficient of Z proteins in the cytoplasm | 1.2 |
| $D_Z^{wall}$ | Diffusion coefficient of Z proteins in cell walls | 0.5 |
| $D_{GFP}^{cyt}$ | Diffusion coefficient of GFP proteins in the cytoplasm | 0.1 |
| $D_{GFP}^{wall}$ | Diffusion coefficient of GFP proteins in cell walls | 0.01 |
| $d_{GFP}$ | Degradation rate of GFP proteins | 0.001 |

